# High-throughput proteomic analysis of FFPE tissue samples facilitates tumor stratification

**DOI:** 10.1101/667394

**Authors:** Yi Zhu, Tobias Weiss, Qiushi Zhang, Rui Sun, Bo Wang, Zhicheng Wu, Qing Zhong, Xiao Yi, Huanhuan Gao, Xue Cai, Guan Ruan, Tiansheng Zhu, Chao Xu, Sai Lou, Xiaoyan Yu, Ludovic Gillet, Peter Blattmann, Karim Saba, Christian D. Fankhauser, Michael B. Schmid, Dorothea Rutishauser, Jelena Ljubicic, Ailsa Christiansen, Christine Fritz, Niels J. Rupp, Cedric Poyet, Elisabeth Rushing, Michael Weller, Patrick Roth, Eugenia Haralambieva, Silvia Hofer, Chen Chen, Wolfram Jochum, Xiaofei Gao, Xiaodong Teng, Lirong Chen, Peter J. Wild, Ruedi Aebersold, Tiannan Guo

## Abstract

Formalin-fixed, paraffin-embedded (FFPE), biobanked tissue samples offer an invaluable resource for clinical and biomarker research. Here we developed a pressure cycling technology (PCT)-SWATH mass spectrometry workflow to analyze FFPE tissue proteomes and applied it to the stratification of prostate cancer (PCa) and diffuse large B-cell lymphoma (DLBCL) samples. We show that the proteome patterns of FFPE PCa tissue samples and their analogous fresh frozen (FF) counterparts have a high degree of similarity and we confirmed multiple proteins consistently regulated in PCa tissues in an independent sample cohort. We further demonstrate temporal stability of proteome patterns from FFPE samples that were stored between one to 15 years in a biobank and show a high degree of the proteome pattern similarity between two types histological region of small FFPE samples, i.e. punched tissue biopsies and thin tissue sections of micrometer thickness, despite the existence of certain degree of biological variations. Applying the method to two independent DLBCL cohorts we identified myeloperoxidase (MPO), a peroxidase enzyme, as a novel prognostic marker. In summary, this study presents a robust proteomic method to analyze bulk and biopsy FFPE tissues and reports the first systematic comparison of proteome maps generated from FFPE and FF samples. Our data demonstrate the practicality and superiority of FFPE over FF samples for proteome in biomarker discovery. Promising biomarker candidates for PCa and DLBCL have been discovered.

## Introduction

Quantitative molecular profiling of phenotypically well annotated clinical sample cohorts using genomic, transcriptomic or metabolomic techniques, followed by the statistical association of molecular and phenotypic data has been a powerful approach for the development of biomarkers, guiding classification, stratification and therapy, particularly with regard to cancer patients ^1,2^. With the increasing robustness, accuracy and throughput of molecular profiling techniques, the need for large, well-annotated sample cohorts has been accentuated over the last few years.

The history of FFPE samples dates back to 1893 ^3^. Most human tissue specimens archived in hospitals for diagnostic purposes are FFPE blocks which have been shown to be stable over time and are usually associated with rich clinical and phenotypic data, including histology, diagnosis, treatment history and response, and outcome. For fresh or rapidly frozen tissue samples such meta data are less frequently available and concerns about molecular stability over time have been raised ^4,5^. FFPE samples have been globally used for DNA, RNA, protein and morphological measurements, and preanalytical factors affecting each type of measurement have been identified ^6^. Besides, various techniques and evaluation studies have been reported for genomic ^7,8^, transcriptomic ^9,10^, proteomic and protein ^11–14^ from FFPE samples.

The preparation of FFPE samples depends on the exposure of the tissue to a range of chemical reactions and conditions. During fixation, formaldehyde reacts with proteins or peptides to form unstable methylol adducts (specified by a C-O bond) which further partially dehydrate to yield active intermediate Schiff bases. These intermediate products subsequently react with basic and aromatic amino acids to form stable and irreversible methylene bridge crosslinks (specified by a C-N bond) ^14,15^, thus modifying the sample proteins. Protein analysis of FFPE tissues using antibodies started in 1991 with the development of the heat-induced antigen retrieval (HIAR) technique for immunohistochemistry (IHC) ^16^. HIAR is based on the notion that heating may unmask epitopes by hydrolysis of methylene cross-links, thus enhancing immunoreactivity. Consequently, the measurement of specific proteins by HIAR has become widely used for diagnostic and prognostic biomarker testing, particularly in cancers ^17^. To extend the analysis of proteins to a proteomic scale, a number of different methods have been used to retrieve proteins from FFPE samples for mass spectrometric analysis ^12,18–24^. They include high pressure ^23^ or pressure cycling technology ^24^ and the available methods have been recently reviewed ^13,14^. These studies have shown that FFPE samples can, in principle, be analyzed by mass spectrometry based proteomic methods. However, the proteome maps of FFPE tissues and their analogue FF tissues from clinical cohorts and their respective stability over time have not been rigorously assessed. The concern remains that FFPE samples may harbor greater variation in protein quality than FF samples due to formalin-induced chemical modifications ^25^.

Multiple factors might have contributed to these limitations. First, the generation of clinically meaningful results requires the consistent analysis of sizable sample cohorts. Second, reproducible sample preparation and mass spectrometric analysis that are essential for clinical studies have been difficult to achieve. Few if any published studies on FFPE proteomic analyses have ever attempted to repeat analysis on clinical specimens of a cohort due to the complexity and high cost of the adopted proteomics techniques ^26–28^. Third, the ability to analyze a histological region of small FFPE samples remains challenging. Most published studies analyzed 26,29,30 tissue micrometer sections with each tissue containing multiple histological types ^26,29,30^. Laser capture microdissection has been used to analyze multiple regions of a tissue section; however, experimental complexities preclude application to large-scale analysis. Targeted needle punches from a FFPE tissue block represent a reasonable compromise; however, efficient extraction of proteins from such a small needle biopsy and further proteolytic digestion of the proteins into peptides for mass spectrometric analysis has not been reported yet. Finally, although methods are available to analyze proteins from human and animal FFPE samples ^12^, concerns remain whether the thus extracted proteins reliably reflect their actual abundance pattern in the fresh frozen counterpart and ultimately, fresh samples ^25^.

In this report we re-visited and optimized the acidic and alkaline hydrolysis procedures developed in 1947 ^31^ which are compatible with a detergent-free protocol to recover proteins from small (0.5×0.5×3mm) FFPE tissue punches in a form that is directly compatible with in solution digestion within an hour. The thus treated tissue samples can be directly processed by the PCT method to generate mass spectrometry-ready peptide samples within a few hours ^32–35^. We further investigated whether the thus acquired FFPE proteome map is comparable to its counterpart FF proteome map in prostate tissue samples by applying this workflow to identify promising diagnostic protein biomarkers for PCa patients. We found that the two types of patterns were highly similar and identified strongly overlapping sets of proteins that showed different levels of expression in benign and tumor tissue. Subsequently, the effect of factors such as storage time and FFPE tissue forms to the proteome was further evaluated. There is no significant difference among FFPE proteome patterns with different storage time, while tissue sections were separated from punched tissue biopsies based on principal component analysis (PCA).

Further, a panel of 12 proteins showing great potential for PCa diagnosis was characterized in an independent Chinese prostate cohort and was validated in the Swiss cohort in this study as well as in other two recently reported PCa studies ^36,37^. As a second application, the FFPE PCT-SWATH workflow was employed to identify prognostic biomarkers for DLBCL patients, employing the relevant archived FFPE tissues. MPO was identified as a promising novel prognostic candidate for DLBCL.

## Results

### Establishment of a FFPE PCT-SWATH workflow

We integrated a workflow for the generation of proteome map from FFPE tissue samples in a robust and high-throughput manner. In addition, we showed that the proteome map derived from FFPE samples correlate well with corresponding maps generated from their analogous FF samples, and that the same biomarker panel can be identified from both sample types, even if the samples have been stored for 4-8 years in their respective format. The de-crosslinking of FFPE tissue is based on acidic and alkaline hydrolysis which was developed in 1947 ^31,38^ but has not been reported in proteomics research applications yet. Here we integrated the classical decrosslinking method with PCT-assisted protein extraction and digestion, and SWATH-MS ^32,39^ method to establish a detergent free FFPE PCT-SWATH workflow. To achieve the desired overall performance profile, protocols for the chemical extraction of proteins from FFPE tissue, liquid chromatography (LC), SWATH-MS, and data analysis were optimized and integrated.

#### Chemical extraction of proteins from FFPE tissue punches

A detergent-free and fast hydrolysis protocol for preparing MS-ready peptides from FFPE tissue punch samples mimicking needle biopsies (width < 1mm, length ~2-3mm; dry mass weight about 300~400μg) (**Fig. 1a**) was optimized. The method consists of i) an acidic hydrolysis step (0.1% formic acid) to achieve C-O hydrolysis of protein methylol products (**Fig. 1b**), ii) a step of heat and base induced hydrolysis to reverse the C-N methylene crosslinks (**Fig. 1b, 1c**) and iii) extraction and digestion of proteins from the thus pre-treated punches by PCT (**Fig. 1c**). The FFPE tissue biopsies used for the protocol establishment were from a sample pool of 48 replicate tissue biopsies extracted from a resected prostate of the ProCOC cohort ^40,41^. We optimized the acidic and alkaline hydrolysis steps by varying the respective treatment times. Acidic hydrolysis was achieved concurrent with the complete rehydration of FFPE tissue punches by replacing water with 0.1% formic acid. Preliminary UV-spectroscopy results showed that the release of methylol groups began saturated in 30 min. As to the alkaline hydrolysis, the effects of the different tested conditions were evaluated by determining the peptide yield as well as the number and type of peptides and proteins identified from each sample by SWATH-MS (**Fig. 1d** and **Fig. 1e**). At this step, 30 min boiling of the FFPE punch with 0.1 M Tris-HCl (pH 10.0) at 95°C led to the highest peptide yield with the greatest number of identified peptides. As shown in **Fig. 1f**, we generated on average of about 60 μg peptide mass per milligram FFPE tissue sample (dry mass with wax). The yield was comparable to our previous investigations of fresh frozen tissues (wet tissue) ^32,33,42,43^.

**Figure 1.**
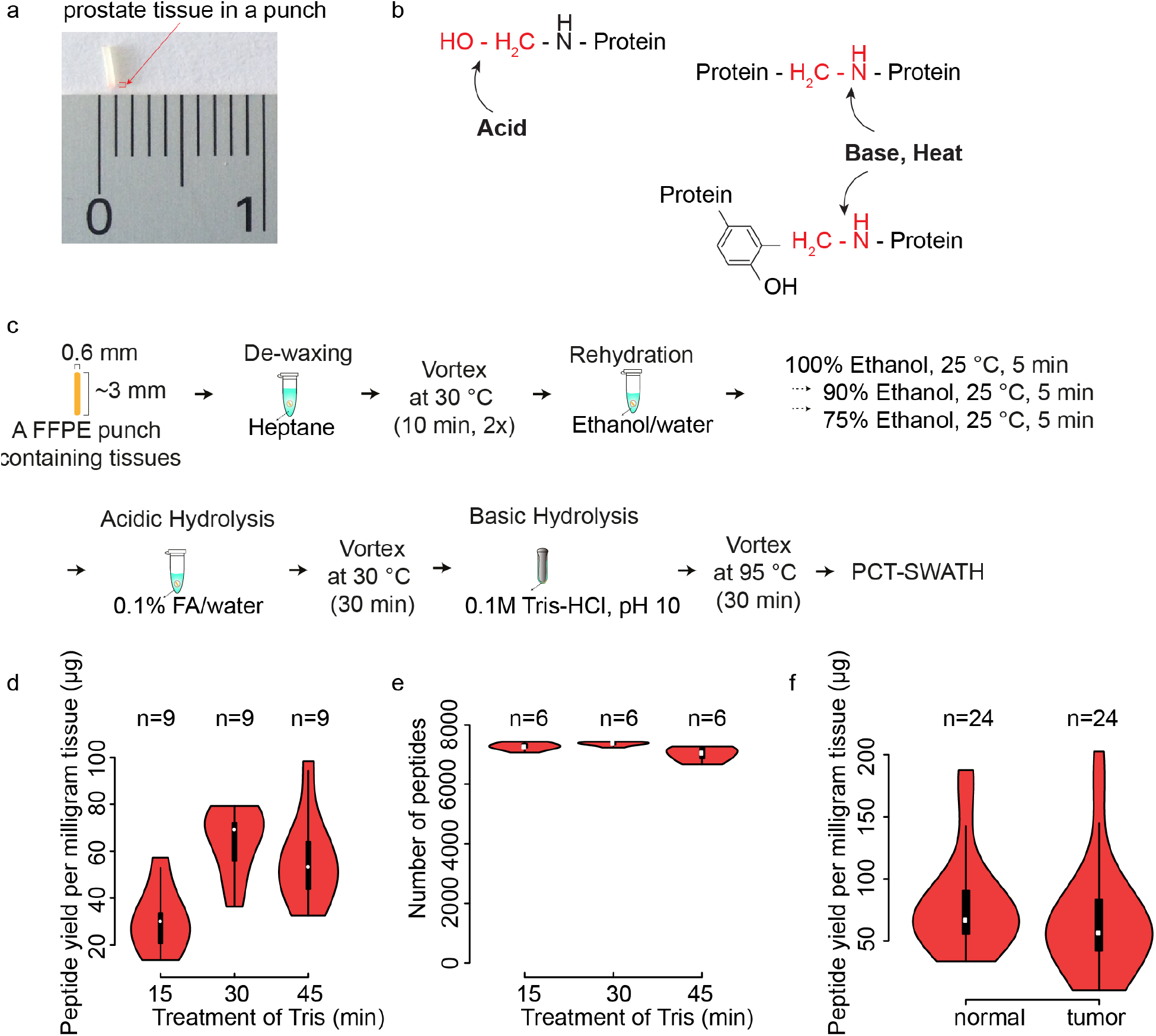
FFPE PCT-SWATH protocol and performance. (**a**) Prostate FFPE tissue in a punch. (**b**) Acid, base and heat treatment to reverse crosslinks. (**c**) Schematic protocol of FFPE PCT-SWATH. (**d**) Peptide yield per milligram FFPE tissue with different Tris-HCl (pH 10.0) boiling time. (**e**) Number of peptides identified by the peptides prepared with different Tris-HCl boiling time. (**f**) Yield of peptides from 48 prostate tissue samples.

#### Optimization of LC and SWATH-MS

We assessed the combined effects of LC gradient length (30 min, 45 min and 60 min) and SWATH window configuration (eight configurations, ranging from 20 to 93 variable windows) on sample throughput, proteome depth and reproducibility. Each window scheme assessed was based on equal segmentation of precursor ion signals over the entire mass range. The peptides used for the optimization of LC and SWATH settings were randomly selected from 9 peptide samples obtained from FFPE tissues processed with the 30 min alkaline hydrolysis protocol described above (**Fig. 1e**). Altogether, we compared 24 LC-SWATH conditions in duplicate (**Supplementary Fig. 1a-1c**). The results showed that 48 variable SWATH windows achieved the highest number of peptide and protein identifications. We observed a trade-off between the gradient length and proteome coverage. The 30 min LC gradient resulted in a 19% lower number of peptide identifications and 8% fewer protein identifications, compared to the 60 min LC gradient (**Supplementary Fig. 1b and Supplementary Fig. 1c**).

### Comparison of FFPE and FF tissue proteome maps

To investigate whether the obtained FFPE proteome maps were comparable to their FF counterparts, we performed proteomic analyses of corresponding FFPE and FF counterpart tissue samples of 24 PCa patients with radical prostatectomy from the ProCOC cohort ^40,41^. Sections of tissue samples from the same resected prostates have been stored for four to eight years in the form of FFPE or FF, respectively, prior to proteomic analysis (**Fig. 2a**, **Supplementary Table 1**). Only the index tumor with the highest Gleason score and the largest diameter was selected for analysis. For non-tumorous tissue, benign prostatic tissue with minimal stromal component was chosen.

With respect to FFPE tissues, three replicate punches were processed via the FFPE PCT-SWATH workflow for each sample and combined for PCT-SWATH analysis. The size of each tissue biopsy was around 0.6×0.6×3mm, weighing approximately 300μg including wax. In total, approximately 900μg of dry mass weight was available per FFPE sample. For the FF cohort, one tissue punch of approximately 1mm^3^ size and wet weight of about 800μg per sample was processed by PCT-SWATH. Altogether, 48 FFPE tissue samples (benign and tumor) were processed into peptide samples that were analyzed by SWATH-MS in technical duplicates. **Fig. 1f** shows that the samples produced on average about 60μg injection-ready peptide mass per milligram of tissue sample. The yield is consistent with previous reports for FF tissues ^32,33,42^. The CV values of peptide yield were 49% and 65% for benign and tumorous tissues respectively, slightly higher than the corresponding figures reported previously for FF tissues ^33,42^. The difference in peptide yield is likely caused by inaccurate estimation of FFPE tissue weight due to variable wax content and the heterogeneity of human prostate tissues.

The setting of 30 min LC integrated with 48 variable SWATH windows was adopted for protein measurement to compare FFPE and FF tissue proteome maps in this study. Two technical replicates for each tissue digest were analyzed using SWATH-MS, referred to in the following as PCF dataset. The resulting SWATH-MS data from all 96 FFPE and FF samples were compared by their total ion current and the number and type of peptides as well as proteins that could be identified and quantified.

We first compared the raw ion intensity signals over chromatographic time (total ion chromatogram, TIC) at both the MS1 and MS2 levels. We found that the TIC, normalized for total injected peptide mass, was on average 15% higher for FF than for FFPE samples (**Supplementary Fig. 2**). The observed small discrepancy of normalized MS1 intensity values is likely due to incomplete acidic and alkaline hydrolysis of cross-links, resulting in the generation of partially hydrolyzed methylene bridges, which contribute to the absorbance in the range from 260-280nm on the spectroscopy. The modification by formalin could also lead to ion suppression. The root cause for lower specific TIC was not further investigated because the effect was minor and the contour of the TIC for FF and FFPE samples were very similar, suggesting that comparable peptide populations were generated from both sample types (**Supplementary Fig. 2**).

Next, we used the SWATH-MS fragment ion maps to compare the number and type of peptides and proteins that could be identified from FF and FFPE samples, and their respective quantities. We used the OpenSWATH ^44^ software tool and a spectral library built from prostate tissue, consisting of 70,981 peptide precursors from 6,686 SwissProt proteins, to search the acquired fragment ion maps. Altogether, we obtained quantitative data for 3,030 SwissProt proteins inferred from 18,129 proteotypic peptides. The median technical CV analyses were 14.9% and 17.5% for FF and FFPE samples, respectively. Overall median CV was 16.2% (**Fig. 2b**, **Supplementary Fig. 3**). We further compared the overall proteomic variation for different tissue types including benign and tumorous FFPE versus FF samples, and found no significant discrepancy (**Supplementary Fig. 4**). We then compared the peptide precursors and proteins detected in each paired FFPE and FF sample (**Supplementary Fig. 5**) and found that peptides as well as proteins were consistently quantified in both tissue types with relatively high Pearson correlation. The overall correlation between FFPE and FF samples reached a Pearson correlation of 0.91 (**Fig. 2c**) with a median normalization of the data based on protein abundance. With unsupervised clustering, the proteome map from FFPE samples was mixed with FF samples (**Supplementary Fig. 6a**), further supporting the notion that the data generated from FFPE samples are comparable with those of FF samples. Curated MS signals by the viewer function of the DIA-expert software for a representative peptide which was quantified across all 224 SWATH runs are shown in **Supplementary Fig. 6b**. We further compared the raw signals, quantity of peptide precursors and proteins in samples stored for different periods of time and observed no significant impact of storage time (**Supplementary Fig. 7**).

Overall the data show that a highly consistent and significant fraction of the whole proteome, consisting of 3,030 SwissProt proteins, could be reproducibly identified from equivalent FF and FFPE samples, even after 8 years storage. Furthermore, the quantitative information generated from matched FFPE and FF sample pairs were comparable.

**Figure 2.**
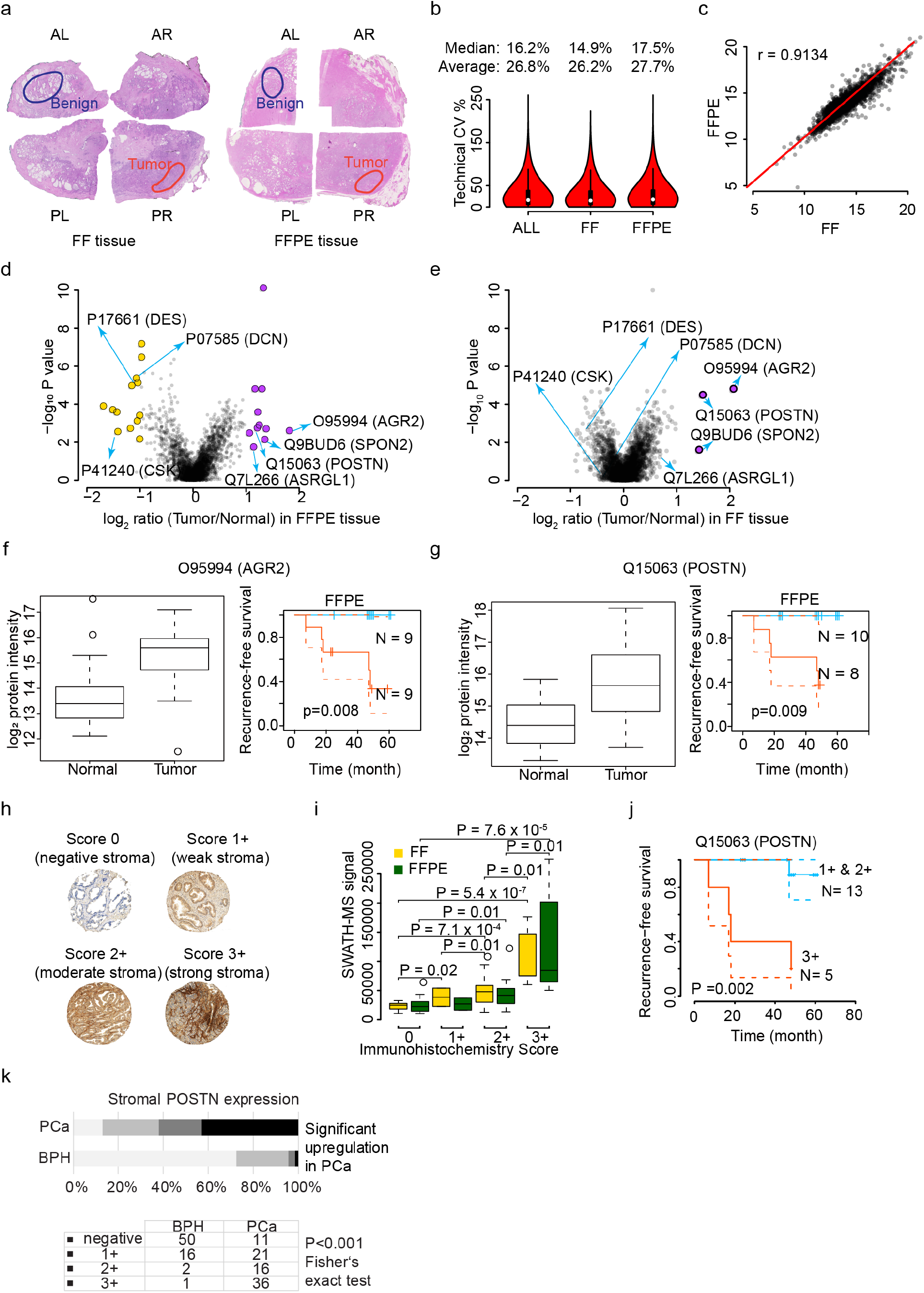
Comparison of FF and FFPE tissues in a patient cohort. (**a**) Benign and tumorous samples were punched from prostate tissue stored since resection as FF and FFPE. The hematoxylin & eosin staining of FF and FFPE tissue from Patient No. 2 in the ProCOC cohort is shown here. AL, anterior left; AR, anterior right, PL, posterior left; PR, posterior right. (**b**) Overall technical CV of FFPE and FF samples at peptide level. (**c**) Comparison of median protein abundance in FF (x-axis) versus FFPE (y-axis) samples. Each dot denotes one protein identified in this sample cohort. Volcano plots show proteins with significant abundance difference between tumor and benign tissue in FFPE (**d**) and FF (**e**) samples from the PCF data set. Proteins showing an abundance difference of fold change (FC) ≥2 and with P value ≤ 0.05 between groups were considered significant. Boxplots and Kaplan-Meier plots show expression of AGR2 (**f**) and POSTN (**g**) in benign and tumorous FF and FFPE samples. (**h**) TMAs of FFPE samples matching those analyzed by mass spectrometry were constructed and stained with an antibody against POSTN. The intensity of stromal POSTN immunoreactivity was scored semi-quantitatively by assigning four scores (0, 1+, 2+, 3+) to each sample. Graphs depict examples of stromal staining. Diameter of each tissue core was 0.6mm. (**i**) Comparison of POSTN expression as measured by immunohistochemistry, and the results from PCT-SWATH in FF and FFPE samples. Statistically significant differences between groups were calculated using two-sided Student *t* test. (**j**) Kaplan–Meier biochemical recurrence-free survival plots of prostatectomy patients stratified by stromal POSTN immunoreactivity in PCa. (**k**) POSTN staining of a TMA.

### Systematic evaluation of the effect of FFPE tissue storage format and duration on proteome maps

We next evaluated the robustness of proteome maps obtained from FFPE tissue stored in different formats and for different periods of time. We procured FFPE tissue samples from three BPH patients from China (termed as ‘PCZC’ cohort). For each patient, we collected both tissue sections (5 μm thickness) and punched tissue biopsies (1×1×0.5 mm). For each sample format we analyzed three biological replicates. The samples had been archived for different periods of time, specifically for 1 year, 5 years, 10 years and 15 years, respectively. Altogether, 72 tissue samples were processed, and 72 SWATH files were acquired with a 90 min LC gradient in a TripleTOF 5600+ mass spectrometer coupled to an Eksigent micro-flow system.

We reproducibly quantified 3,040 SwissProt proteins in both tissue punches and sections in this data set. By comparing the protein abundance distribution of these common proteins, we found that the proteome maps of the two FFPE formats showed a high degree of similarity, with a Pearson correlation coefficient of 0.95 (**Fig. 3a**). The mean Pearson correlation coefficient of all 72 samples among their own biological replicates was 0.858, showing that the samples were of high similarity at the whole proteome level (**Fig. 3b**). Unsupervised cluster analysis of all 3,040 proteins also showed consistent distribution of protein abundance among all 72 samples (**Fig. 3c**). We further grouped the 72 samples into eight groups according to sample format and storage time, and investigated the biological variation of nine samples (three patients, each with three biological replicates) in each group. The average CV slightly varied between tissue micrometer sections and punches across the time span of 15 years (**Fig. 3d**). Further, tissue micrometer sections were found to be different from punches (**Fig. 3e**), probably due to that fact that tissue micrometer sections cover more diverse tissue regions and therefore contain higher degree of the spatial heterogeneity ^43^. However, these differences only affected a small portion of proteins. The duration of FFPE storage did not impact on our proteomic measurement, further reinforcing the stability of FFPE proteome and the robustness of our protocol (**Fig. 3f**).

**Figure 3.**
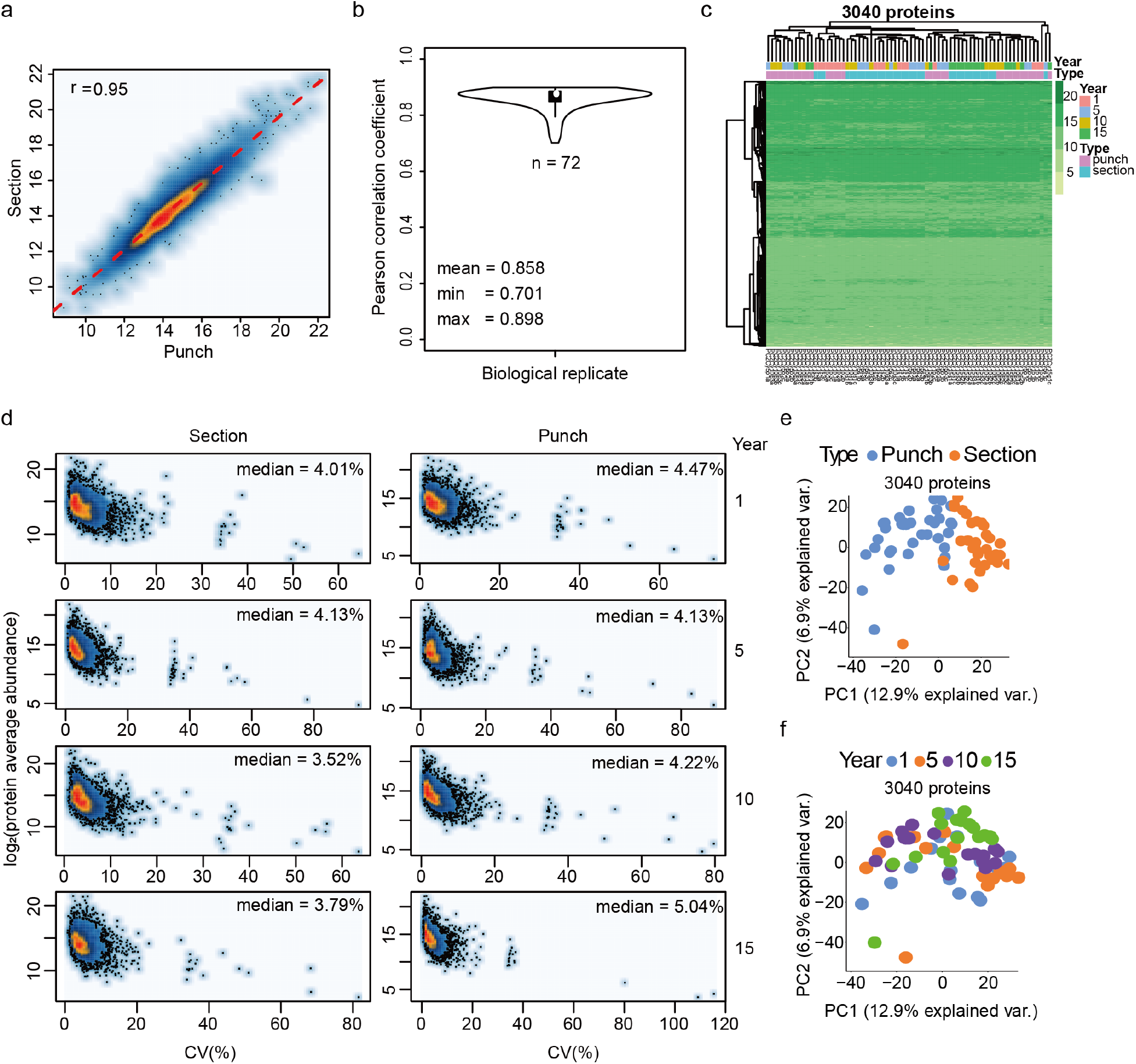
Evaluation of FFPE tissue storage forms and duration. (**a**) Pearson correlation of protein abundance between FFPE micrometer sections and punches. (**b**) Average Pearson correlation coefficient of all 72 samples among three biological replicates. The ‘pairwise.complete.obs’ method was employed to calculate the COR value to avoid the influence of NA. (**c**) The protein abundance distribution of all 3,040 SwissProt proteins across all 72 samples with different tissue types and storage time. (**d**) CV plots for each sample type (section/punch) with different storage time (1 yr, 5 yrs, 10 yrs and 15 yrs). (**e**) PCA analysis of the effect of tissue types. (**f**) PCA analysis of the effect of storage time.

### Identification of a subset of proteins with comparable abundance patterns in prostate FFPE and FF punches

Next, we asked whether proteins distinguishing benign and tumorous prostate tissue could be consistently detected in both FFPE and FF samples. We observed differential expression of multiple proteins between benign and tumorous tissues in both FFPE and FF samples in the PCF cohort.

We first determined proteins that were significantly differentially regulated between the FF tumor and benign samples. We computed the median fold-change of tumor-to-benign tissue samples and the P values for each protein in the 24 patients for FF tissue samples. By setting a fold-change (FC) cutoff of 2 and a P value cutoff at 0.05, only three proteins were significantly up-regulated in tumor compared to benign tissue. These were Q15063 (POSTN), O95994 (AGR2) and Q9BUD6 (SPON2). Remarkably, these three proteins are all promising biomarker candidates. POSTN is an extracellular matrix protein involved in cell development and adhesion. We have previously reported its upregulation in high grade and advanced stage PCa patients ^45^, which is consistent with an independent report of its positive prognostic value in PCa^46^, and with a study of its positive correlation with the aggressiveness of PCa ^47^. AGR2 is a secreted adenocarcinoma-associated antigen. The mRNA level of AGR2 was found higher in cancerous tissue in 42 paired PCa samples, but it was not associated with survival in the cohort ^48^. In addition, the protein expression level of AGR2 was found increased in cancerous tissue in 31 out of 58 PCa cases by IHC immunolabeling ^48^, a result that was consistent with two independent cohorts ^49^. In this study, the expression of AGR2 was found to be about four times higher in tumor compared to benign tissue and, remarkably, it’s abundance level positively correlated with survival (P = 0.008). SPON2 is a secreted extracellular matrix protein. In a previous study, it was detected as an abundant protein in serum samples of 286 PCa patients compared to 68 healthy controls ^50^. In particular, it was found with significantly higher expression levels in PCa patients with a Gleason score of 7-8 and in PCa patients with metastases ^50,51^. In our FF data set, SPON2 was found to be expressed 2-4 times higher in malignant compared to corresponding benign tissue samples.

We then analyzed the SWATH data acquired for FFPE samples in the same way and found 24 proteins with significantly different abundance between tumor and benign groups. The results from the FFPE cohort recapitulated the pattern of the three proteins with increased tumor abundance identified in the FF cohort. The consistency of the detected changes for these proteins is remarkable given the intra-tumor heterogeneity, expected differences between the FFPE and FF proteomes and the fact that the FFPE and FF samples were from different regions of the tumors. In addition to these three proteins, a further eight proteins were detected at increased abundance in FFPE tumor compared to benign tissue and thirteen proteins were detected at lower abundance in the tumor vs. benign samples (**Fig. 1d**).

To verify whether the findings from our SWATH data set of ProCOC ^40^ patients are consistent with IHC reports, we analyzed a tissue microarray from 18 patients which were also part of the cohort analyzed by PCT-SWATH. Representative staining images of POSTN are shown in **Fig. 2h**. We scored the staining patterns into four grades (0, 1+, 2+, and 3+), and compared the results with the SWATH signals of the corresponding FF and FFPE samples. We analyzed the statistical significance by pair-wise comparison of groups using Students’ *t*-test, and single factor ANOVA (**Fig. 2i**, **Supplementary Table 2**). We did not observe any significant difference between the data from FF samples and from FFPE at the level of the mass spectrometry data. Remarkably, the ANOVA analyses revealed significant correlation between IHC and both FF (P value 3.3×10^−6^) and FFPE (P value 6.7×10^−5^) SWATH data. Taken together, via the IHC results an orthogonal technique confirmed the similarity of POSTN abundance patterns of POSTN detected in FFPE and FF samples by mass spectrometry. Nevertheless, the difference among the IHC groups 0, 1+ and 2+ appeared mostly insignificant at the SWATH level. The prognostic role of POSTN was further confirmed in the survival analysis based on the TMA data (**Fig. 2j**). In an independent Swiss TMA cohort, we also observed significantly higher abundance of POSTN (P value < 0.001 by Fisher’s exact test, **Fig. 2k**) in tumor vs, benign tissue.

We then checked the functions and applications of the 24 proteins significantly regulated proteins in the FFPE sub-cohort based on literature mining. Here we discussed some of them which had been studied and reported extensively. Q7L266 (ASRGL1) was found to be significantly upregulated in FFPE tumor samples, whereas the quantitative difference in FF samples was not significant. The full name of ASRGL1 is isoaspartyl peptidase/l-asparaginase protein, which is an enzyme involved in the production of l-aspartate. ASRGL1 was overexpressed in PCa and regarded as the potential diagnostic and therapeutic target ^52^. Among the 13 down-regulated proteins identified in FFPE cohort, desmin (DES, P17661) is a known marker protein for prostate smooth muscle ^53^. The decreased abundance of DES in tumor tissue is consistent with the replacement of smooth muscle tissue by malignant cells. We also found that c-Src tyrosine kinase (CSK, P41240), a regulator of SRC kinase ^54^, was found to be down-regulated in tumor tissue. With respect to decorin (DCN, P07585), a proteoglycan in the tumor microenvironment, our data for the first time report its down-regulation in association with PCa prognosis. This observation is in line with a previous mouse-based functional study reporting that DCN specifically inhibits EGFR and AR phosphorylation, leading to suppressed AR nuclear translocation and inhibition of PSA production ^55^. While most protein changes were detected in both tissue types, the FFPE samples exposed the protein regulation with better statistical power (**Fig. 2**). POSTN was detected to be significantly upregulated in both FF and FFPE tumor samples in this cohort. CSK and DCN were only significant in the FFPE cohort, indicating the FFPE proteomes analyzed by our method are more robust.

Furthermore, by integrating the seven proteins (POSTN, AGR2, SPON2, ASRGL1, DES, CSK, and DCN) discussed above, we achieved an AUC of 0.983 for FF samples and 0.977 for FFPE samples, respectively (**Fig. 4a**) for the separation of tumor and benign tissue. Our data again demonstrated the consistency of FFPE and FF proteome maps acquired by the PCT-SWATH workflow and the ability to identify differentially abundant proteins from either sample type. Further, the data shows that the observed abundance differences were attenuated in in FF samples compared to their FFPE counterparts. This could be due to gradual protein degradation during long-term storage in the frozen state.

**Figure 4.**
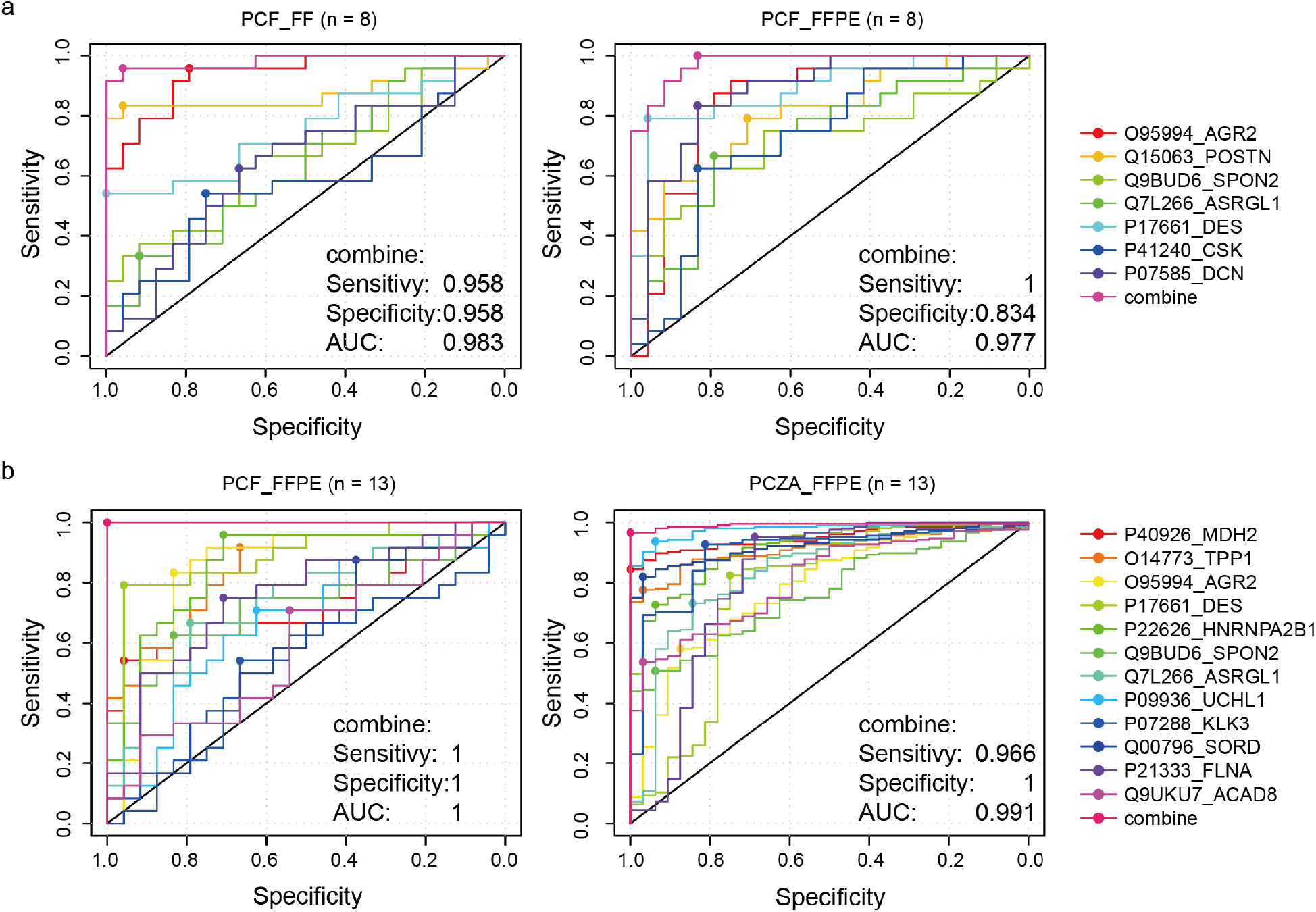
Receiver Operating Curves (ROC) analysis for the diagnostic power of protein signature panels. (**a**) ROC of seven regulated proteins and the combined model using logistic regression in FF (left) and FFPE (right) samples from the PCF cohort. (**b**) ROC of 12 regulated proteins and the combined model using logistic regression in PCF-FFPE (left) and PCZA (right) samples. Area Under Curve (AUC) values are shown.

### Integrative protein signature for stratifying PCa

We applied the same workflow as above to another cohort from China (in the following termed as ‘PCZA’ cohort) to stratify PCa. The PCZA cohort contains samples from 58 PCa patients and 10 benign prostate hyperplasia (BPH) patients that have been stored as FFPE samples for up to 2 years. Three punches for each sample were analyzed. Of these samples, we randomly selected 33 for technical replicates (**Supplementary Table 3**). Altogether, 237 SWATH files were acquired. To cover more proteins, we adopted an extended 120 min LC gradient in a TripleTOF 5600+ coupled to an Eksigent micro-flow system. The resulting data were processed as described above. From 4,144 SwissProt proteins quantified with high degree of reproducibility (**Supplementary Fig. 8**) in this data set, we identified 241 up-regulated proteins and 89 down-regulated proteins (adjusted P value cutoff 0.05, FC cutoff 2) (**Supplementary Table 3, Supplementary Fig. 8**). We performed ingenuity pathway analysis (IPA) ^56^ of these significantly regulated proteins between PCa and BPH groups and found that five top upstream regulator pathways were enriched from these proteins (**Supplementary Table 4**). MYCN, MYC, TCR regulator pathways were activated while sirolimus and 5-fluorouracil regulator pathways were inhibited (**Supplementary Table 4**). 16 cellular networks were enriched from these proteins via IPA analysis (**Supplementary Table 4**).

PCZA and PCF datasets shared seven common regulated proteins in prostate tumor tissues, which are O14773 (TPP1), O95994 (AGR2), P22626 (HNRNPA2B1), P40926 (MDH2), Q9BUD6 (SPON2), P17661 (DES), Q7L266 (ASRGL1), as shown in **Fig. 4b**. Besides, other significantly regulated proteins, including P07288 (KLK3), Q00796 (SORD), P21333 (FLNA), P09936 (UCHL1), and Q9UKU7 (ACAD8), were also identified to show diverse functions by IPA analysis (**Fig. 4b**, **Supplementary Table 4**). These proteins were enriched in nine networks by IPA analysis, as shown in **Supplementary Fig. 9**. The relative abundance of these proteins in both cohorts was calculated and the regulation pattern of them in two cohorts was consistent with each other, as was shown in **Fig. 5**. The regulation of these proteins between tumor and benign tissues was much more significant in PCZA cohort as was demonstrated by P values. PCZA consists two groups of 58 PCa patients and 10 BPH patients, while PCF contains the tumor/benign pair of tissue from 24 PCa patients. 3,030 proteins were quantified from PCF cohort by the 30 min LC plus 48-variable-window scheme from 224 Swath files in Zurich, while 4,144 proteins were quantified in PCZA cohort by the 120 min LC plus 48-variable-window scheme from 237 Swath files in Hangzhou, both using AB Sciex TripleTOF 5600+. The two cohorts shared 2846 proteins in common, accounting for 93.9% of the PCF whole proteome. Then we calculated the Pearson correlation of PCF and PCZA FFPE tumor proteomes with the r value 0.514, reflecting the existence of certain degree of biological variations between the Swiss and Chinese cohorts (**Supplementary Fig. 8**). By loosening the threshold for significantly regulated proteins in PCF cohort, more proteins would be distinguished out to be deregulated between tumor and benign conditions, as was shown in **Fig. 5**.

**Figure 5.**
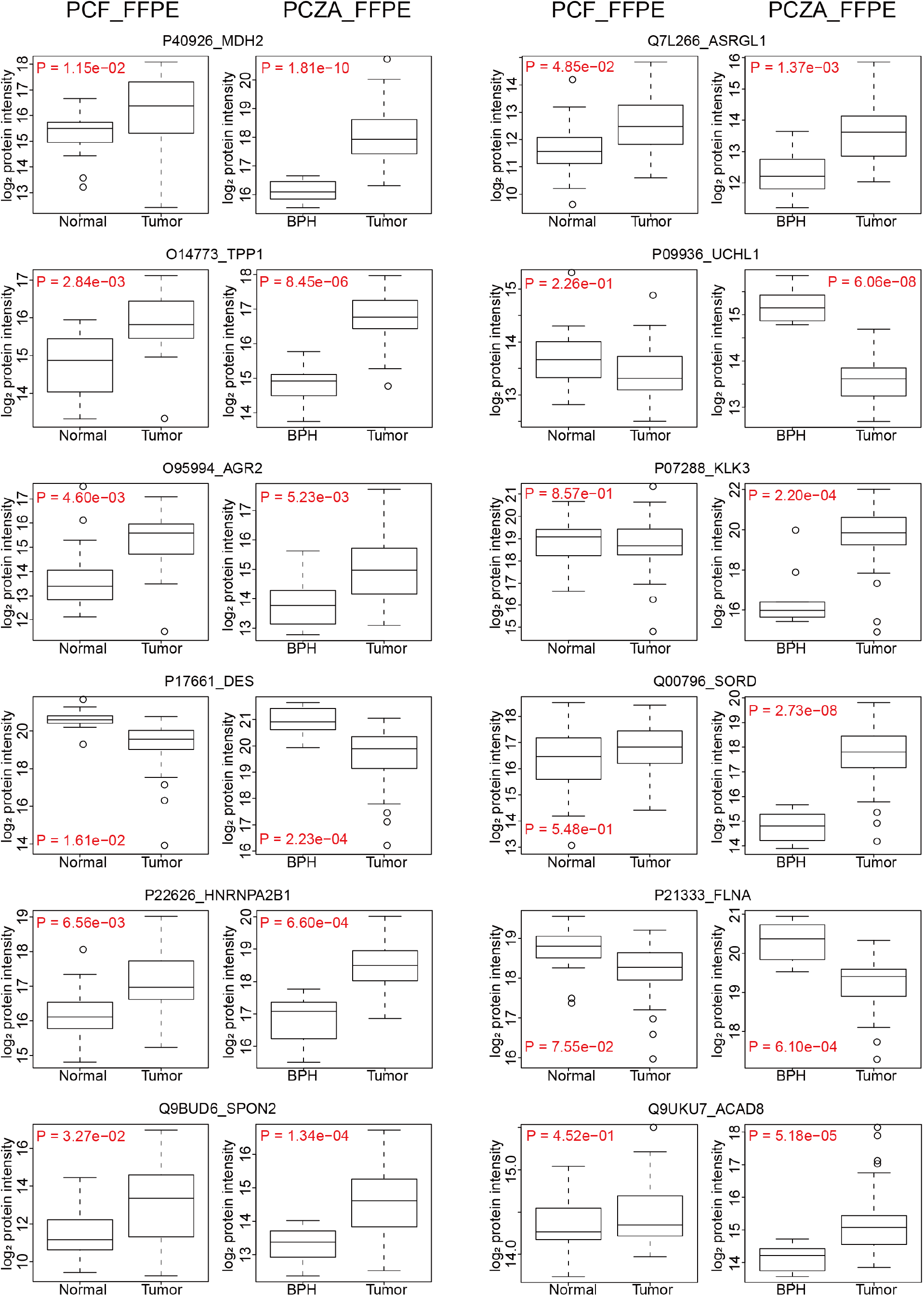
Relative abundance of the twelve proteins in paired normal and tumor prostate samples in PCF cohort and BPH/tumor samples in PCZA cohort, respectively. FLNA, UCHL1, and DES were downregulated in tumor tissues, while others were upregulated.

KLK3 is the prostate-specific antigen (PSA), a serum marker for PCa. Sorbitol dehydrogenase (SORD) converts sorbitol to fructose. SORD is part of the polyol pathway that plays an important role in sperm motility. SORD is regulated by androgens in the human prostate, and reported to be positively associated with Gleason scoring and serum PSA concentrations ^57^. Our data show that both KLK3 and SORD were significantly overexpressed in PCa tissues. Both Filamin-A(FLNA) and -B (FLNB) were proposed as protein panel signatures for diagnosis of PCa ^58,59^. FLNA was found to be downregulated in PCa tissues. UCHL1 is a ubiquitin-protein hydrolase involved in the processing of ubiquitin precursors. Our data show significant suppression of UCHL1 in tumor tissues, in agreement with a previous report, further consolidating its value in PCa biology^60^.

Tripeptidyl-peptidase 1 (TPP1) was found to be upregulated. TPP1 is a primary protector of telomere DNA and has been reported to be an effective anticancer target for about 90% of human tumors that are telomerase-positive ^61^. Heterogeneous nuclear ribonucleoproteins (HNRNPs) associate with nascent pre-mRNAs, and package them into HNRNP particles in a sequence-dependent way. HNRNP particles serve to condense and stabilize the transcripts and minimize tangling and knotting. The splicing factor HNRNPA1 has been reported to contribute to enzalutamide resistance by promoting AR-V7 ^62^. In this study, HNRNPA2B1 was found to be a novel upregulated protein probably modulating splicing in PCa cells. Malate dehydrogenase 2 (MDH2) was also up-regulated in prostate tumor tissues in both PCF and PCZA cohort in this study. MDH2 is a mitochondrial enzyme that catalyzes the NAD/NADH-dependent, reversible oxidation of malate to oxaloacetate. Interestingly, a very recent report on integrative proteomics in PCa uncovers two metabolic shifts in the citric acid cycle (TCA cycle) during PCa development and progression, among which MDH2 is a component. Increased MDH2 expression in PCa correlated with an increase in mRNA levels, and it is further upregulated in CRPC samples ^36^. Together, these data suggest that development of MDH2 inhibition could be of great benefit against progressed PCa. Besides, ACAD8, the acyl-CoA dehydrogenase family member 8, was detected to be upregulated in tumor tissues in this study. It has been reported to be a potential prognosis biomarker indicating the outcome of prostate tumors ^63^.

We further applied the 12-protein panel to both the Swiss and Chinese PCa cohorts, to evaluate the sensitivity and specificity in diagnosis of PCa. These proteins and their ROC curves using the PCF and PCZA FFPE data sets are shown in **Fig. 4b**. They exhibited high AUC values. Integrative models demonstrated AUC values of 1 in the FFPE samples of the PCF cohort. In the independent PCZA cohort, the AUC reached 0.991. An independent FFPE cohort from a different country therefore confirmed the diagnostic significance of these novel proteins in PCa. Taken together, these findings demonstrate that our proteomic methodology is robust and has the capacity to uncover new diagnostic protein biomarkers for PCa.

Subsequently we identified differentially expressed proteins distinguishing patient groups classified by Gleason scores. In this study, 24 PCa patients from the PCF cohort and 58 PCa patients from PCZA cohort were classified into three groups according to their tumor grades as reflected by Gleason, namely, low (L), intermediate (M), and high stage (H) (**Supplementary Table 5**). ANOVA analysis was employed to compare proteomes among three stages to identify protein candidates that distinguish different stages of cancer progression (P value < 0.05). 216 proteins and 373 proteins were detected significantly regulated in the PCF cohort and the PCZA cohort, respectively (**Supplementary Table 5**), with 23 proteins overlapping. PCA analysis (**Supplementary Fig. 10**) demonstrated clear separation of L and H grades, however, it was challenging to distinguish M from L and H grades, consistent with the pathological nature of the samples, indicating that proteome acquired by our method well preserved the granularity of the FFPE tissue samples.

### Prognostic markers for diffuse large B-cell lymphoma (DLBCL)

Having established that the PCT-SWATH method was applicable to analyze prostate FFPE samples and to consistently distinguish malignant and benign samples in two independent sample cohorts, we next asked whether the method could stratify other types of tumors based on overall survival. We procured 41 patients with DLBCL (in the following termed as ‘WLYM’ cohort) from the University Hospital Zurich to investigate prognostic markers. DLBCL is a disease with relatively poor prognosis and includes different subtypes, *i.e.* lymphomas residing exclusively in the brain, known as primary central nervous system lymphomas (PCNSL) and extracerebral DLBCL (eDLBCL). Another distinct entity, intravascular lymphoma (IVL), is a rare type confined to the lumina of blood vessels (there is only one IVL patient in WLYM cohort, **Supplementary Fig. 11**). About 70% cases of eDLBCL are curable, however, the median survival of patients with PCNSL is only about 30 months in contemporary clinical trials ^64^.

To identify prognostic proteins for DLBCL, two to three FFPE punches were analyzed for each of the 41 DLBCL tumors (**Supplementary Table 6, Supplementary Fig. 11, 12**). Altogether, we acquired 113 SWATH maps using a 60 min LC gradient, and a TripleTOF 6600 mass spectrometer. We quantified 5,769 SwissProt proteins in all samples (**Supplementary Table 6**). The technical reproducibility for a representative sample is shown in **Supplementary Fig. 11d**. 91 proteins were detected to be significantly up-regulated and 6 proteins were detected to be down-regulated in the PCNSL tumors compared to eDLBCL tumors (**Supplementary Fig. 11e, Supplementary Table 6**). Of these, 20 proteins were suspected to be contaminants from brain tissue based on their brain tissue expression annotation in the DAVID database and the human protein atlas (**Supplementary Table 6**) ^65^. 17 proteins were further selected from the remaining 77 proteins according to their applications in biomarker and drug target studies as revealed by IPA analysis ^56^ (**Supplementary Table 6**). Their relative abundance of these proteins in both eDLBCL and PCNSL groups is shown in **Supplementary Fig. 13**.

ROC analyses based on these seventeen proteins in both eDLBCL and PCNSL patient samples from WLYM cohort were performed. Two proteins including glial fibrillary acidic protein (P14136, GFAP) and zeta chain of T cell receptor associated protein kinase 70 (P43403, ZAP70) exhibited high AUC values (**Fig. 6a**) to differentiate eDLBCL and PCNSL subtypes of DLBCL. GFAP is a class-III intermediate filament and a cell-specific marker that distinguishes astrocytes from other glial cells during the development of the central nervous system. We found that GFAP is a novel upregulated marker in PCNSL. ZAP70 is a tyrosine kinase that is essential for initiation of T cell antigen receptor signaling. ZAP70 deficiency is associated with Immunodeficiency 48 that is a form of severe immunodeficiency characterized by a selective absence of CD8+ T-cells ^66^. Here we found that ZAP70 was upregulated in the eDLBCL subtype compared with PCNSL, indicating the role of ZAP70 in immunological processes during the progress of the disease.

To further investigate the prognostic value of the proteins identified above, we procured a second cohort of 52 eDLBCL patients from China (in the following termed as ‘ZLYM’ cohort), and performed FFPE PCT-SWATH analysis using a TripleTOF 5600+ coupled to an Eksigent micro-flow LC system (**Supplementary Table 7**). Two biological replicates were analyzed for each patient. Here we quantified 6,266 proteotypic SwissProt proteins in 52 micro-sectioned tissue samples from these DLBCL patients in technical duplicate. 16 out of 17 proteins identified in the WLYM cohort described above were also identified in the ZLYM cohort. Survival analysis of the 16 proteins in both groups of eDLBCL patients (WLYM and ZLYM) was further performed through Kaplan-Meier plot. The result showed that besides ZAP70, five additional proteins namely crystallin alpha B (P02511, CRYAB), Golgi membrane protein 1 (Q8NBJ4, GOLM1), myeloperoxidase (P05164, MPO), microtubule associated protein 1A (P78559, MAP1A) and ubiquitin C-terminal hydrolase L1 (P09936, UCHL1), were found to show consistent trend in predicting the survival outcome in both WLYM and ZLYM eDLBCL patient cohorts, although the P values in most cases are not very significant due to the small size of the cohorts that were available for this rare disease (**Fig. 6b**). CRYAB has the function of preventing aggregation of various proteins under a wide range of stress conditions. GOLM1 is highly expressed in colon, prostate, trachea and stomach. Our study identified them as novel biomarkers for eDLBCL patients.

MPO is a lysosomal protein known as expressed in azurophilic granules (primary lysosomes) of normal myelomonocytic cells which is released into the extracellular space during degranulation. MPO functions as part of the host defense system of polymorphonuclear leukocytes. It is responsible for microbicidal activity against a wide range of organisms. MPO has been reported to be related to myeloperoxidase deficiency (MPOD) that is characterized by decreased myeloperoxidase activity in neutrophils and monocytes that results in disseminated candidiasis ^67^. MAP1 is a structural protein involved in the filamentous cross-bridging between microtubules and other skeletal elements. MAP1A/B are neuron specific microtubules ^68^. MAP1S has been reported to interact with mitochondrion-associated leucine-rich PPR-motif containing protein (LRPPRC) that interacts with the mitophagy initiator and Parkinson disease-related protein Parkin ^69^. UCHL1 gene mutations are involved in Parkinson disease 5 (PARK5) that is characterized by a complex neurodegenerative disorder with manifestations ranging from typical Parkinson disease to dementia with Lewy bodies ^70^. As discussed above, UCHL1 is also a tumor suppressor in a broad range of cancers including PCa. eDLBCL patients with lower expression level of MPO, MAP1, UCHL1 and ZAP70 were found to have higher survival rate in this study.

Higher expression of MPO in eDLBCL patients was associated with worse survival, as was shown in Kaplan-Meier plot (**Fig. 6b**). IHC staining of MPO in DLBCL tumors from two patients in WLYM cohort confirmed the presence of MPO-positive regions (**Fig. 6c**). Detection of increased abundance of MPO in eDLBCL group compared to the PCNSL group might indicate the presence of coagulative necrosis with penetration of MPO^+^ granulocytes in the aggressive subset of DLBCLs ^71^. Taken together, the data suggest that MPO is a robust prognostic marker for DLBCL patients. This also supports the robustness of this proteomic methodology, even if independent sample cohorts are studied in different laboratories and instruments. The data from punches from the WLYM cohort matched well with the sectioned samples from the ZLYM cohort.

**Figure 6.**
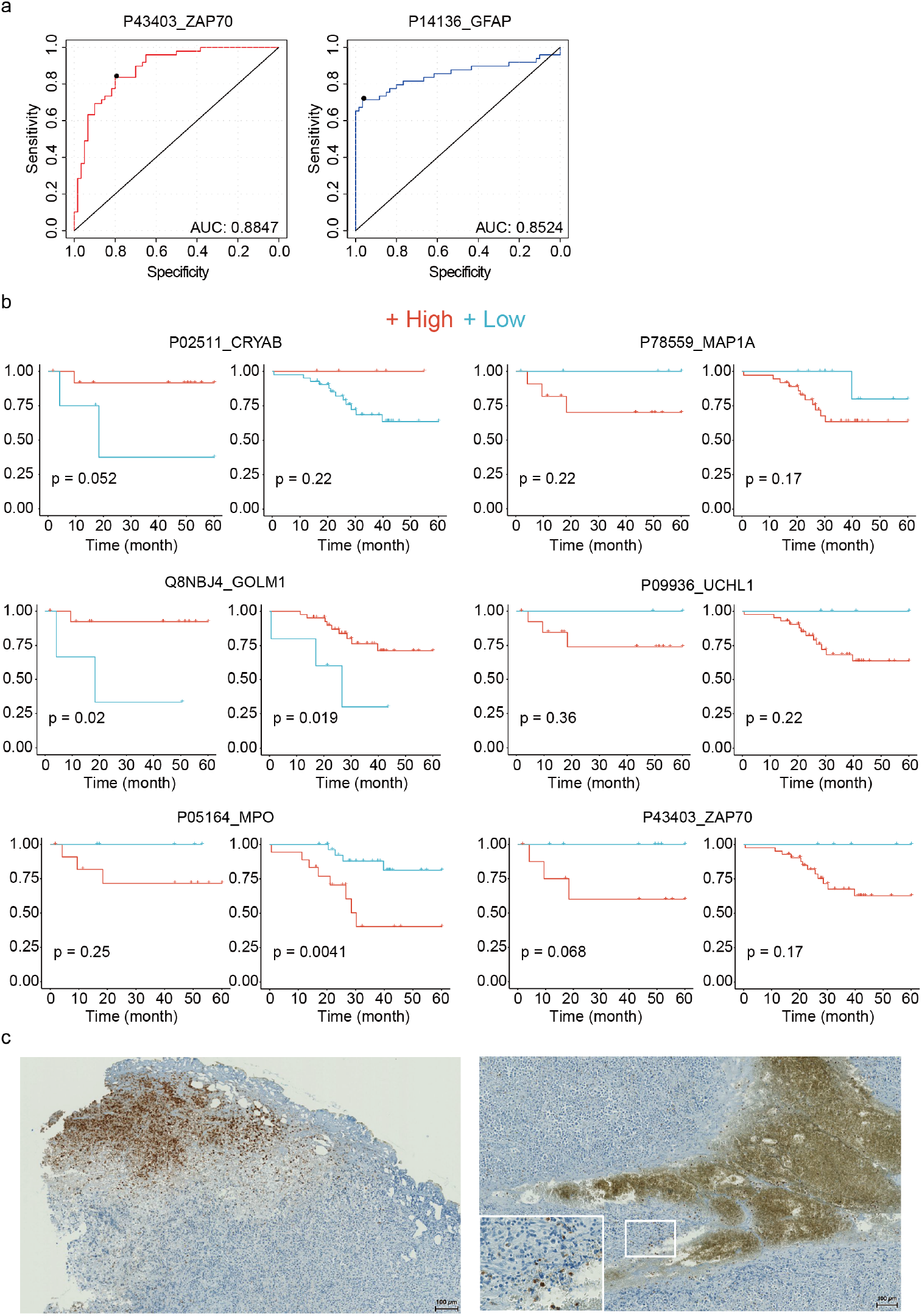
(a) ROC analysis for the diagnostic power of GFAP and ZAP70 to distinguish eDLBCL and PCNSL subtypes in WLYM cohort. (**b**) Survival analysis of a six-protein panel for eDLBCL patients in both WLYM and ZLYM cohorts. (**c**) Representative IHC staining of MPO in two eDLBCL patients in the Zurich WLYM cohort.

## Discussion

Most archived tissues in pathology collections exist as FFPE samples, representing a rich resource for clinical research. Over the past decade, MS-based shotgun proteomics has been used to analyze proteins from FFPE samples ^12–14,18–24,72^. However, the concern remains that FFPE samples may harbor greater variation in protein quality than FF samples due to formalin-induced chemical modifications ^25^. Ostasiewicz *et al.* performed a comparison of FFPE and FF mouse liver tissues and found similar protein pattern^12^. However, this was not confirmed in human tissues. Recently, Piehowski, *et al.* analyzed 60 FFPE ovarian cancer samples with the storage from 7 to 32 years using TMT 10-plex isobaric labelling method coupled with shotgun proteomics approach and reported no significant proteome expression difference in terms of age and storage time ^30^. This is an informative study investigating the clinical value of FFPE samples, however, the practicality, robustness and reproducibility of FFPE proteomics, in terms of sample preparation and LC-MS analyses, has not been rigorously established. Procurement of a suitable cohort sample for rigorous comparison of FFPE and FF samples is critical for validating the practicality.

In this study, based on the ProCOC cohort ^40^ which allowed access to prostate tissue samples from adjacent sections of the same resected tissue was stored in both FFPE and FF format with the storage over 4 to 8 years, we performed rigorous proteomic comparison between them. PCT-SWATH analysis of 224 PCa FFPE and FF samples facilitated a rigorous comparison in a clinical scenario in this study. Regarding to the storage factors that might affect the whole proteome, comparison of proteome maps of FFPE samples stored for 8 years and for 4 years did not show significant pattern differences (**Supplementary Fig. 6**). A further overall investigation of FFPE sample proteome maps storing from 1 year to 15 years in an independent cohort (PCZC) did not show significant pattern differences either (**Fig. 3**). Besides, proteome maps from two types of FFPE tissue forms (sections vs. punches) are generally similar, however, they could be separated from each other by PCA analysis (**Fig. 3**).

Since proteins in FFPE tissue are extensively and substantially modified by formalin ^14,73^, one would not expect complete recovery of the entire proteome, and quantitatively identical recovery of every peptide in various samples. Previous studies have investigated this issue in depth ^74,75,76^. Indeed, we observed a slight global difference in TIC between comparable FFPE and FF proteomes (**Supplementary Fig. 2**). However, we also show that these differences do not distort the proteome patterns to a degree that would preclude their use for tissue classification, suggesting that the slight differences observed between FF and FFPE tissue samples are smaller or comparable to other preanalytical factors ^77^. This observation is significant because frequently, longitudinal sample collections that are invaluable for biomarker discovery are stored in FFPE format. Remarkably, despite a number of potential confounding factors, we successfully identified the same protein biomarker candidates from matching FFPE and FF samples in the ProCOC cohort ^40^, even though the FFPE and FF samples were actually from different, albeit consistently scored sections in these prostate samples. To the best of our knowledge ^12–14,18–24,72^, this is the first study in which the proteome of FFPE and FF has been rigorously compared in a clinical scenario.

Regardless of the variable formalin fixation processes of tissue specimens, reproducible sample preparation and LC-MS analysis are essential for clinical studies. Due to the complexity (dozens to hundreds of fractions for a single sample) and high cost of the lengthy shotgun proteomic workflow (hundreds to thousands of MS analyses for a single cohort), few published studies on FFPE/FF proteomic analyses have ever attempted to repeat analysis on clinical specimens of a cohort ^26^. A rapid and robust methodology for quantitatively measuring proteomes of FFPE tissue specimens at low-cost and in a high-throughput manner are in great need.

In this study, we identified a panel of twelve-protein biomarker candidates including KLK3, SORD, AGR2, SPON2, MDH2, ACAD8, TPP1, DES, HNRNPA2B1, ASRGL1, UCHL1 and FLNA as differentially abundant between tumor and benign tissues from two independent PCa cohorts, PCF and PCZA (**Fig. 5**). With this panel, the malignant tissue can be separated from benign prostate tissue with an AUC higher than 0.9 in both PCF and PCZA sample cohorts (**Fig. 4b**). To evaluate the quantitative PCa proteome maps generated in this study, as well as to investigate the biological differences among different PCa cohorts from different countries, we compared our PCF and PCZA proteomes with the two representative PCa proteomes generated by Iglesias-Gato et al ^37^, and Latonen et al ^36^, respectively. 3030 proteins were quantified from PCF cohort, and 4,144 proteins from PCZA cohort in this study. Iglesias-Gato et al used the Super-SILAC labeling plus multi-fractionation integrated with shotgun MS method, to profile proteotypes of 28 prostate tumors (Gleason score 6–9) FFPE samples and neighboring nonmalignant FFPE tissue in eight cases (sections of 10μm thickness and 25 mm^2^ area), and quantified 1,216 proteins from over 9000 protein identifications ^37^. Latonen et al reported high-throughput SWATH-MS proteotyping of fresh clinical tissue samples (five 5 μm slices for each sample) of 10 BPH patients, 17 untreated PCa patients and 11 CRPC. In PCa vs BPH, they quantified 3,394 proteins, which is comparable with our results regarding to quantified protein number. Moreover, ACO2 and MDH2, two components in TCA cycle during PCa development and progression were identified ^36^. In our study, the overexpression of MDH2 in PCa tissues in both PCF and PCZA cohort was characterized, which was consistent with Latonen’s report. Venn diagram showed that PCF, PCZA and Latonen cohorts shared 2,277 common proteins in total, representing 67% of the Latonen proteome, as was shown in **Supplementary Fig. 14 and Supplementary Table 8**. PCF, PCZA and Iglesias-Gato cohorts shared 700 proteins in total, representing 57% of the quantified proteome by Iglesias-Gato. Besides, in Iglesias-Gato cohort, five proteins from our 12-protein panel were found to be significantly regulated, which were MDH2, TPP1, UCHL1, FLNA, and ACAD8. In Latonen cohort, six proteins, MDH2, TPP1, AGR2, DES, HNRNPA2B1, and ACAD8 were found to be significantly regulated. Detailed information of protein regulation of the twelve-protein biomarker candidates was shown in **Supplementary Table 8**. The four cohorts revealed common proteins biomarkers and showed good consistence although there were biological differences. Taken together, the presented data not only demonstrate the practicality of using FFPE samples for robust PCa biomarker discovery, more importantly, it also identified a panel of protein biomarker candidates for PCa diagnosis, among which MDH2, TPP1 and ACAD8 were most significant regardless of tissue formats (fresh or FFPE, punch or micrometer section) and patient populations. The overlap of the four proteomes confirmed the technical reliability, robustness and transferability of our FFPE PCT-SWATH pipeline among different studies, cohorts and laboratories from another point of view.

The hereinabove studied PCa cohort offers a rational model to benchmark the similarity of FFPE and FF proteome due to the availability of both types of tissue samples from adjacent regions with relatively high degree of homogeneity. However, PCa patients generally exhibit positive prognosis after prostatectomy. To further explore the generic applicability of the method and to explore the feasibility of identifying prognostic markers in another clinical setting, we analyzed 113 FFPE samples from a cohort of 41 Swiss DLBCL patients from Zurich with up to 125-month follow-up. We further validated the methodology and findings using the independently established FFPE PCT-SWATH platform in China, which comprised 52 Chinese DLBCL patients with up to 100-month follow-up. Importantly, data from the two cohorts confirmed MPO as a promising survival marker (**Fig. 6**). The discovery of MPO as a potential prognostic marker for DLBCL is also supported by the finding that circulating monocytes and neutrophils are reported to be independent prognostic factor for DLBCL ^78^. Myeloid cells are presumably MPO-positive and found to suppress T-cell responses. It also indicates the presence of coagulative necrosis.

In conclusion, we demonstrated that FFPE tissue cohorts effectively facilitate biomarker discovery compared to its FF counterpart via the optimized FFPE PCT-SWATH proteomics analysis. We also reported novel promising protein biomarkers for PCa and DLBCLs. This study indicates that historical FFPE tissue samples from biobanks have great potential in biomarker discovery.

## Materials and Methods

### Prostate tissue Specimens

Both FF and FFPE tissues from Zurich were kindly provided by P.J.W. in the form of punches from the Department of Pathology and Molecular Pathology, University Hospital Zurich. Samples were collected within the ProCOC study ^40^, a prospective ongoing biobanking trial led by PJW and CP. The size of a single FF tissue biopsy was about 1mm^3^ (diameter 1 mm; length 1-2 mm; wet weight was about 800 μg). The size of a single FFPE punch is about 0.5×0.5×3mm; and the dry mass weighed about 300 μg including wax (**Fig. 1a**). The Cantonal Ethics Committee Zurich (KEK-ZH) has approved all procedures involving human material, and each PCa patient has signed an informed consent form (KEK-ZH-No. 2008-0040). Patients were followed on a regular basis, every three months during the first year and afterwards at least annually or on an individual basis depending on the disease course. A PSA value of 0.1 ng/ml or higher was defined as biochemical recurrence ^79^.

Prostate FFPE samples from China (PCZA cohort) were procured in the Second Affiliated Hospital of College of Medicine, Zhejiang University with approval from the hospital ethics committee. The size of a single FFPE punch of PCZA cohort is about 1×1×5mm; and the dry mass weighed about 1-1.5 mg including wax. Three punches were collected from each tissue sample.

Prostate FFPE samples from China (PCZC cohort) were procured in the First Affiliated Hospital of College of Medicine, Zhejiang University with approval from the hospital ethics committee. The cohort contained two forms of archived FFPE samples that are micrometer sections (5μm thickness) and tissue biopsy punches (1×1×0.5mm). Three biological replicates were collected for each sample. The samples have been archived for a variety of timespans of 1 year, 5 years, 10 years to 15 years.

The PCa study here was also approved by ethics committee of Westlake University.

### Lymphoma Specimens

FFPE tissues from 41 patients with PCNSL, eDLBCL and IVL who had signed an informed consent and had been treated at the University Hospital Zurich between 2005 and 2014 were collected. Studies were approved by the Institutional Review Board (KEK-StV-Nr.19/08). FFPE punches (each FFPE punch measuring about 0.5×0.6×3mm) were produced in the Department of Pathology, University Hospital Zurich. Two to three punches were collected from each tissue sample.

An independent cohort of 52 DLBCL patients was procured from the First Affiliated Hospital of College of Medicine, Zhejiang University with approval from the hospital ethics committee. Five sections with thickness of 10μm were collected for each tissue sample.

The DLBCL study here was also approved by ethics committee of Westlake University.

### De-paraffinization, Rehydration, and Hydrolysis of FFPE tissues

The FFPE tissue was put into a 2 mL safe-lock Eppendorf tube, and was firstly subjected to the dewaxing step by incubation with 1mL of heptane, and then gently vortexed at 800rpm for 10 min at 30 °C on a thermomixer (cap closed). The dewaxing was repeated once. The sample was then subjected to gradient rehydration steps and gently vortexed at 800 rpm with 1ml 100%, 90%, 75% of ethanol, respectively, each time at 25 °C for 5 min (cap closed). At the third step, each tissue sample was further incubated with 200μl 0.1% formic acid (FA) for 30 min at 30 °C for rehydration and the acidic hydrolysis, by gently vortexing at 800rpm (cap closed). Lastly, the tissue sample was transferred into a PCT-MicroTube (Pressure Biosciences Inc., South Easton, MA), briefly washed with 100 μl of 0.1M Tris-HCl (pH 10.0, freshly prepared) to remove remaining FA, and was then incubated with 15μl of fresh 0.1 M Tris-HCl (pH 10.0), boiled at 95 °C for 30 min, by gently vortexing at 600rpm to undergo a heat and base induced hydrolysis (cap closed).

Once hydrolysis was finished, the PCT-MicroTube with FFPE tissue bathed in Tris-HCl (pH 10.0) was immediately placed on ice for cooling, then 25 μl of lysis buffer (6 M Urea, 2 M thiourea, 5 mM Na2EDTA in 100mM ammonium bicarbonate, pH 8.5) was added to the solution (final pH around 8.8). Both the tissue sample and supernatant from this step were kept for subsequent PCT-assisted tissue lysis and protein digestion.

### PCT-assisted tissue lysis, protein extraction, and protein digestion

Briefly, the FFPE tissue sample was lysed with lysis buffer in a barocycler NEP2320-45k (PressureBioSciences Inc.) at the PCT scheme of 30s high pressure at 45kpsi plus 10s ambient pressure, oscillating for 90 cycles at 30°C. Then the extracted protein solution was reduced and alkylated by incubating with 10mM tris(2-carboxyethyl)phosphine (TCEP) and 20mM iodoacetamide (IAA) at 25°C for 30 min, in darkness, by gently vortexing at 800rpm in a thermomixer. Afterwards, proteins were firstly digested by Lys-C (Wako; enzyme-to-substrate ratio, 1:40) in the barocycler using the PCT scheme of 50s high pressure at 20kpsi plus 10s ambient pressure, oscillating for 45 cycles at 30°C. Then a subsequent tryptic digestion step followed (Progemga; enzyme-to-substrate ratio, 1:20) using the PCT scheme of 50s high pressure at 20kpsi plus 10s ambient pressure, oscillating for 90 cycles at 30°C. Peptide samples were then acidified by TFA prior to C18 desalting. The FF tissue samples were processed as described previously ^32^ with only the change of replacing the normal microcaps with micropestles ^33^.

### SWATH mass spectrometry

All samples were spiked with iRT peptides (Biognosysis) ^80^. 0.6 μg of cleaned peptides (0.3 μg per injection, in technical duplicates) were analyzed by SWATH-MS on a 5600 (Zurich ProCOC cohort) or 6600 TripleTOF mass spectrometer (Zurich DLBCL cohort) connected to a 1D+ Nano LC system (Eksigent, Dublin, CA) ^32,43^. The LC gradient was mixed with buffer A (2% acetonitrile and 0.1% formic acid in HPLC water) and buffer B (2% water and 0.1% formic acid in acetonitrile). The analytical column was home-made (75 μm × 20 cm) using a fused silica PicoTip emitter (New Objective, Woburn, MA, USA) and 3 μm 200 Å Magic C18 AQ resin (Michrom BioResources, Auburn, CA). Peptide samples were separated with a linear gradient of 2% to 35% buffer B over 30 min (the PCF cohort) or 60 min (the WLYM cohort) LC gradient time at a flow rate of 0.3 μl·min^−1^. Ion accumulation time for MS1 was 50 ms and 40 ms for MS2 acquisition, respectively. SWATH window schemes were optimized to 48 variable windows. The instrument was operated in high sensitivity mode.

The PCZA, PCZC and ZLYM cohorts from China were analyzed in a TripleTOF 5600+ coupled to an Eksigent Nano LC 415 (with a 1-10ul/min flowmodule to switch the LC from nano-flow to micro-flow). Composition of mobile phase was the same as that in Zurich lab. Eksigent Analytical column (0.3 x 150 mm C18 ChromXP 3μm) and trap column (0.3 x10 mm, C18 ChromXP 5μm) were used for chromatographic separation. 2μg of peptide samples were separated with a linear gradient of 3% to 25% buffer B over 90 min (the PCZC and ZLYM cohort) or 120 min (the PCZA cohort) LC gradient time at a flow rate of 5 μl·min^−1^. SWATH acquisition method was the same as the method at Zurich lab except a slightly longer MS2 accumulation time of 60 ms.

### SWATH data analysis

We built a SWATH assay library after analyzing unfractionated prostate tissue digests prepared by the PCT method in Data Dependent Acquisition (DDA) mode on a 5600 TripleTOF mass spectrometer over a gradient of 2 hours as previously described ^81^. SWATH data were first analyzed using OpenSWATH (openms 1.10.0) ^81^ as described ^32^. Retention time extraction window was 300 seconds, and m/z extraction was performed with 0.05Da tolerance. Retention time was calibrated using iRT peptides. The sample for each peptide precursor that was identified by OpenSWATH with the lowest m_score was treated as the reference sample for each peptide precursor, and was used as input for DIA-expert analysis (https://github.com/tiannanguo/dia-expert). Briefly, all *b* and *y* fragments for each identified peptide precursor in the Spectrast library were re-analyzed using OpenSwathChromatogramExtractor (openms 1.10.0) for all samples. A reference sample was selected for each peptide precursor based on the m_score from OpenSWATH analysis described above. For reference sample, peptide fragments forming good peak shape were refined in all samples. Peptide precursors with less than four good peak-forming fragments were excluded. Each sample, except the reference sample, in the sample set was pair-wise compared with the reference sample at fragment level, and the median proportion of all fragments was used for quantification of the peptide precursor in a sample. The MS2-level total ion chromatogram for each SWATH window was used to normalize the peak group area. Peptide precursors that were quantified in technical duplicates with a fold-change value equal or higher than two were excluded. The most reliable peptide precursor from a protein, *i.e.* best flier peptide, was selected to represent the abundance of a protein because we found that inclusion of poorly responded peptide precursors negatively influenced to the quantitative accuracy, and that for high abundance proteins with multiple peptides, the best flier peptide selected by the DIA-expert was representative and exhibited the lowest number of missing values. All codes are provided in Github.

### Tissue Microarray and Immunohistochemistry

The Ethics Committee of the Kanton St. Gallen, Switzerland approved all procedures involving human materials used in this St. Gallen TMA, and each patient signed an informed consent. The construction of TMA and IHC procedures have been was described previously ^43^. The POSTN antibody was from abcam (ab14041). The MPO antibody was from NeoMarkers / Lab Vision Corporation (RB-373-A1).

### Statistical analysis

All plots were produced with R. Violin plots were made using the R package vioplot. Pearson’s correlation was used to compute the correlation coefficient. Two-tailed paired Student’s *t*-test was employed to compute probability in Volcano plots. Kaplan–Meier estimators were used for RFS analysis. Point-wise 95% confidence bands were computed for the whole range of time values. Differences between survival estimates were evaluated by the log-rank test.

### Data deposition

The PCF data are deposited in PRIDE ^82^. Project accession: PXD004691. The PCZA data are deposited in iProX (IPX0001355000). The ZLYM data are deposited in iProX (IPX0001354001). All the data will be publicly released upon publication.

## Supporting information

STable1

STable2

STable3

STable4

STable5

STable6

STable7

STable8

## Acknowledgements

This work was supported by the SystemsX.ch project PhosphoNet PPM (to R.A. and P.J.W.) and TPdF 2013/134 (to P.B.), the Swiss National Science Foundation (grant no. 3100A0-688 107679 to R.A.), the Foundation for Scientific Research at the University of Zurich (to P.J.W.), the European Research Council (grant no. ERC-2008-AdG 233226 and ERC-2014-AdG670821 to R.A.) and the Westlake Startup to T.G‥ T.W. acknowledges the Swiss National Science Foundation (323630_164218) and the Swiss Cancer Research Foundation (MD-PhD-3558-06-2015). We thank Andreas Beyer for his critical input to this manuscript, and Patrick Pedrioli for help in data management. We thank the Aebersold group, in particular Moritz Heusel, for invaluable discussion. Members of the tissue biobank of the University Hospital Zurich are thanked for excellent support in providing tissue punches, specifically Peter Schraml, Susanne Dettwiler, Fabiola Prutek, and Peggy Tzscheetzsch. Andre Fritsche, Norbert Wey, Christiane Mittmann, Monika Bieri, Andre Wethmar and Annette Bohnert are thanked for their technical support. The study nurse Alexandra Veloudios is thanked for help with the ProCOC study.

## Author contributions

Y.Z., T.G., T.W., P.J.W. and R.A. designed the project. Y.Z. conceived and developed the hydrolysis protocol and the whole FFPE PCT-SWATH workflow. P.J.W., Q.Z., A.C., K.S., D. R., J.L., C.D.F., M.B.S., C.F., N.J.R., C.P. and W.J. procured the Zurich PCa cohort. T.W., E. R., M.W., P.R., E.H., and S.H. procured the Zurich DLBCL cohort. X.Y. and L.C. procured the Chinese PCa cohort. S.L., B.W. and X.G. procured the Chinese DLBCL cohort. Y.Z. and T.G. optimized the LC-SWATH-MS. Y.Z, T.W., R.S., X.Y., P.B., L.G., C.C., and T.G. performed the PCT-SWATH analysis. Y.Z., T.G., T.W., Q.Z., Z.W., T.Z., C.X. analyzed the data. Y.Z., T.G. T.W. and R.A. wrote the manuscript with inputs from all co-authors. R.A., P.J.W. and T.G. supported and supervised the project.

