## supplemental figures for "High-throughput proteomic analysis of FFPE tissue samples facilitates tumor stratification"

**Supplementary Figure 1**

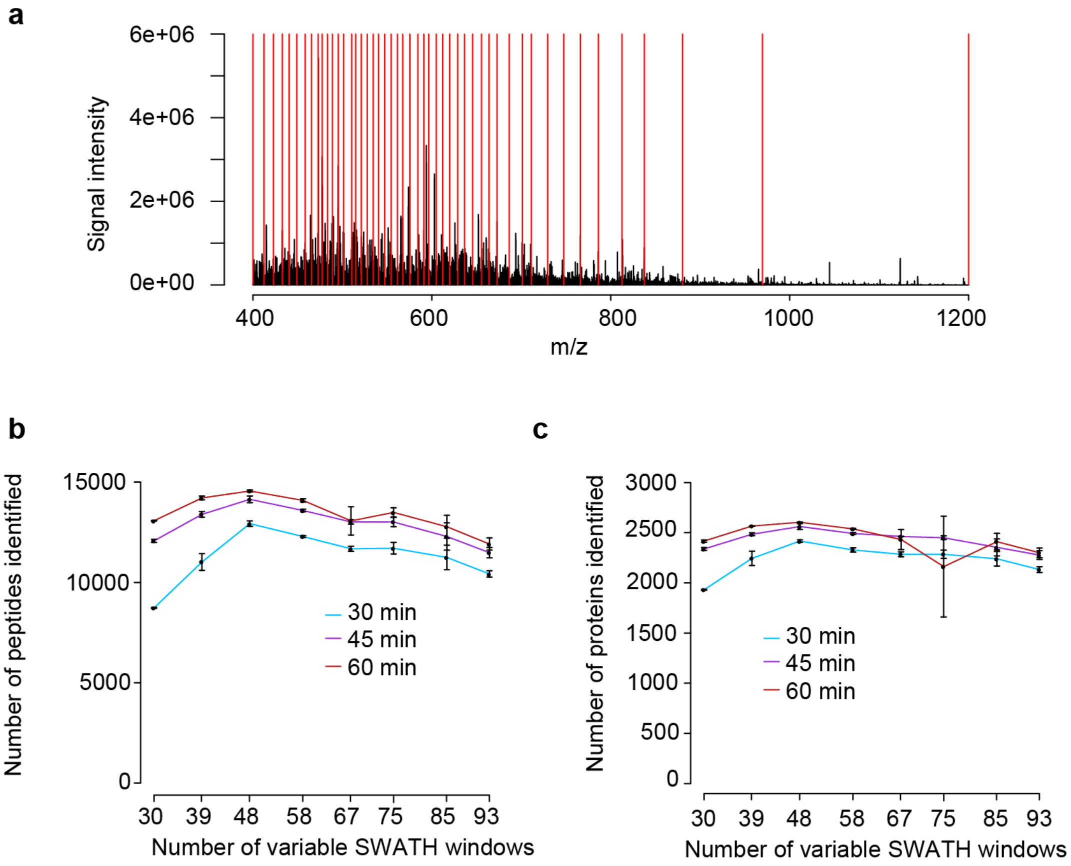


**Supplementary Figure 1. Optimization of LC and SWATH-MS.** (**a**) The design of variable SWATH windows based on quantiles of peptide intensity as exemplified by the 48-variable-window scheme. Comparisons of peptide identification (**b**) and protein identification (**c**) for the 24 LC-SWATH conditions are shown.

#### Supplementary Figure 2

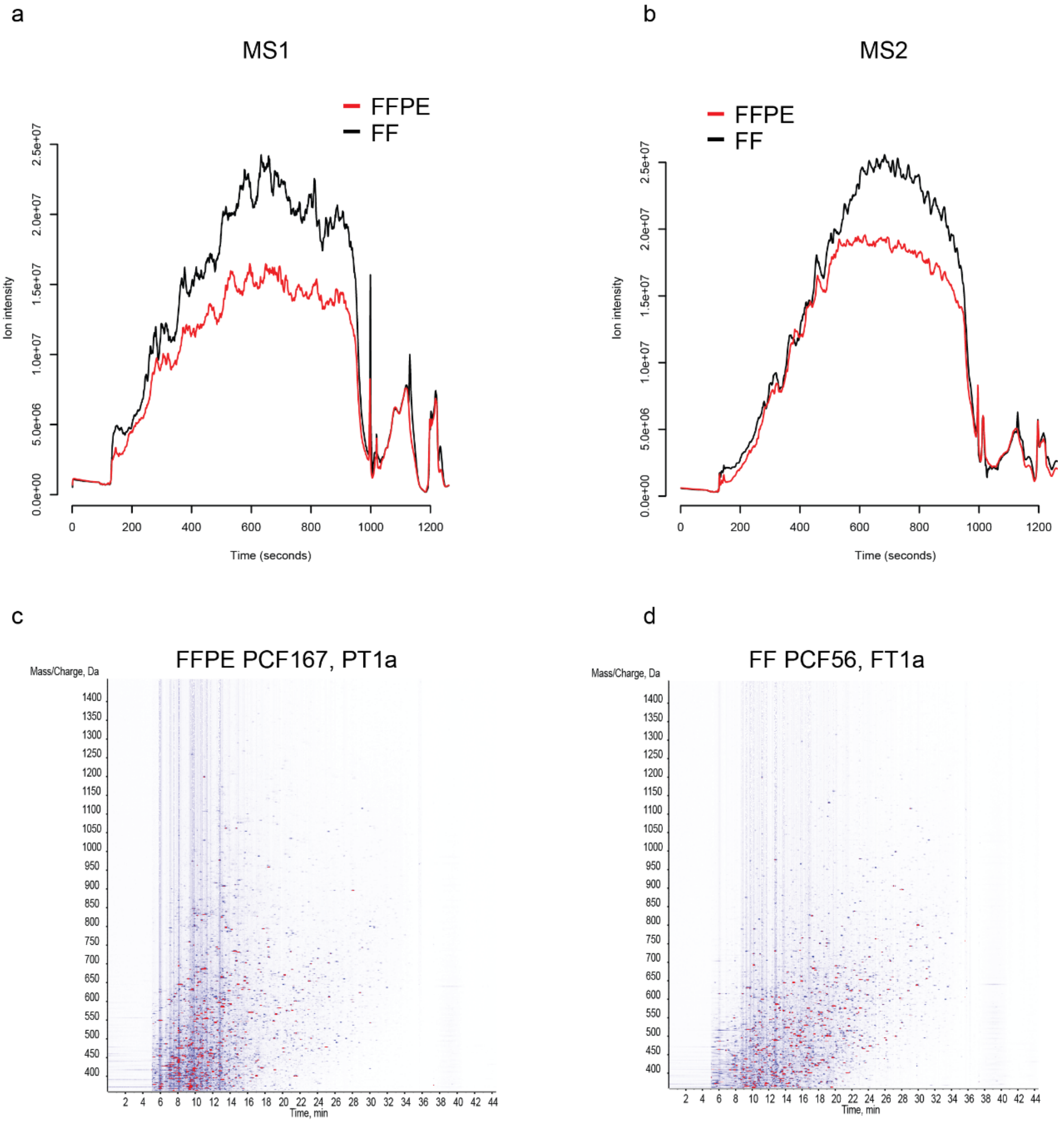


**Supplementary Figure 2. Comparison of the raw SWATH signals from FFPE and FF samples in the PCF dataset.** Total ion chromatogram averaged over all FFPE and FF samples at MS1 level (**a**) and MS2 level (**b**). Representative ion contour map generated from software PeakView for a FFPE **(c)** and the counterpart FF **(d)** sample. The colors indicate the MS signals. Signals higher than 1% of the max intensity are shown as red.

#### Supplementary Figure 3

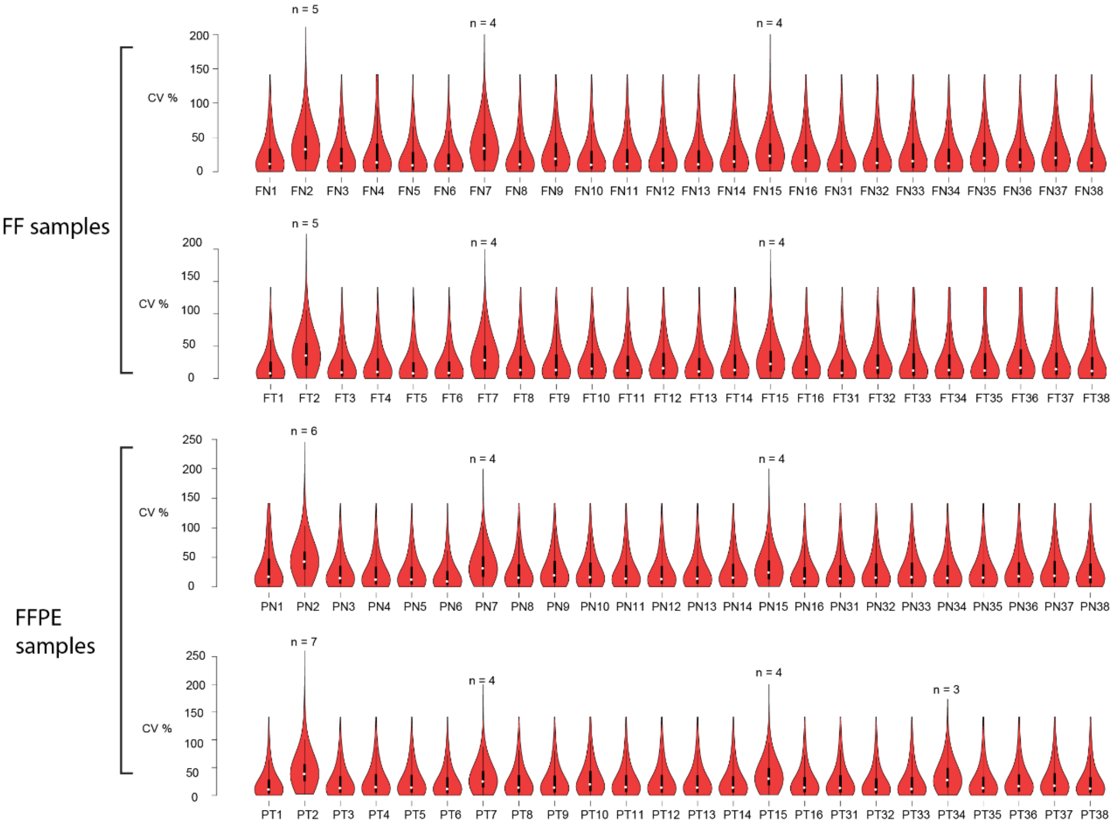


**Supplementary Figure 3. Reproducibility of peptide quantification in the in the PCF dataset.** The distribution of CV values for each of the 192 tissue samples processed in duplicates are shown. Thirteen randomly selected samples were processed in more than two replicates as indicated in the figure. FN: FF tissue, non-tumorous; FT: FF tissue, tumorous; PN: FFPE tissue, non-tumorous; PT: FFPE tissue, tumorous.

**Supplementary Figure 4**

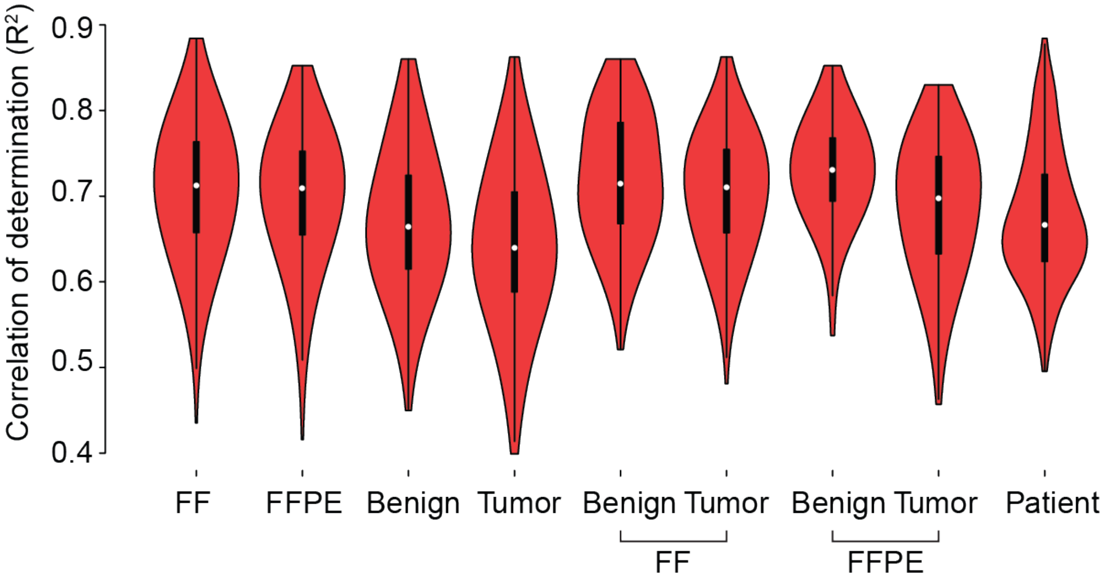


**Supplementary Figure 4. Comparison of overall proteomic variation in different tissue types in the PCF data set.** The correlation of determination ($R^2$) values between pairs of proteome maps for each tissue type were computed and their distributions are shown in a violin plot.

**Supplementary Figure 5a**

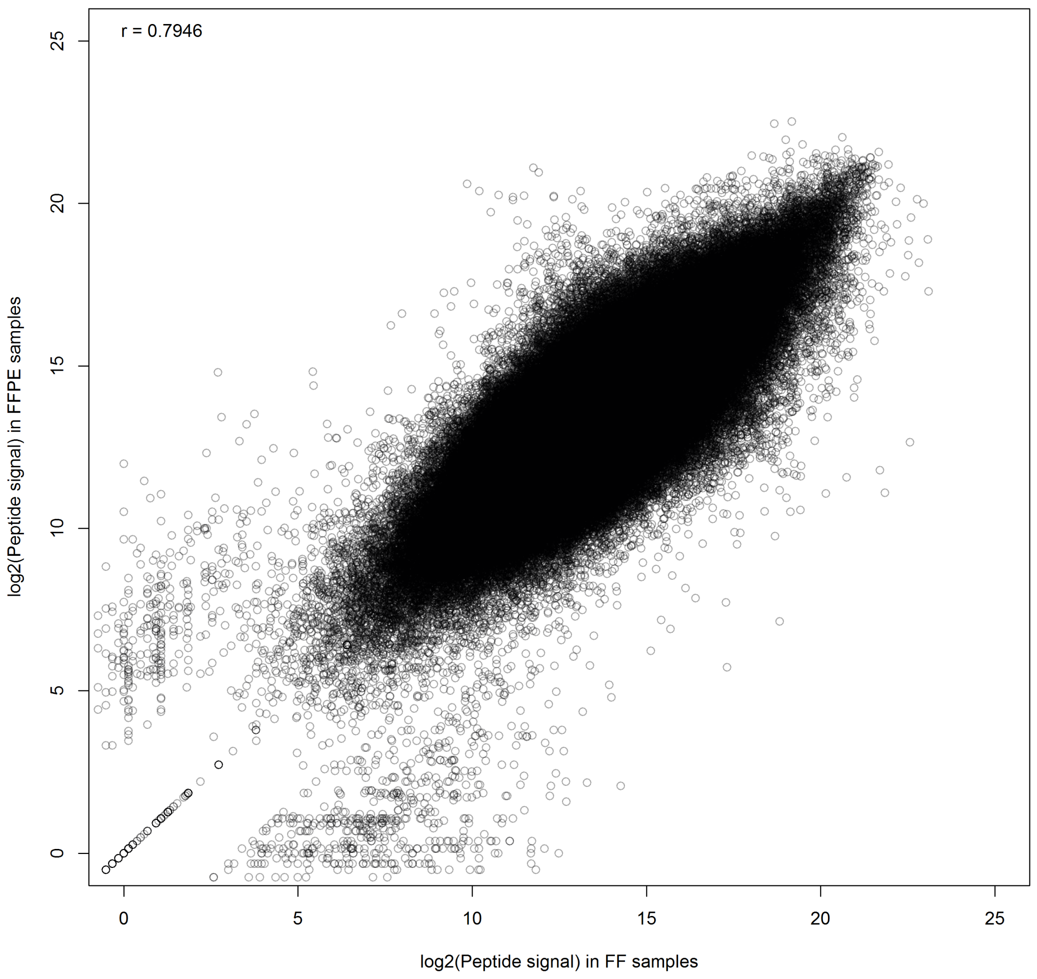


**Supplementary Figure 5a. Comparison of peptide precursor signal in FF and FFPE samples in the PCF dataset.** Each dot indicates a peptide precursor that was quantified in both FF (x-axis) and FFPE (y-axis) samples from the same PCa tissue. Its log2-transformed signals are mapped in the two axes. The overall Pearson correlation is 0.79, indicating that the peptides were detected in both tissue types and their signals correlated. Nevertheless, there was also some variation in the signal, which is not surprising considering the fact that the FF and FFPE tissue samples were a from adjacent sections of the same tissue.

**Supplementary Figure 5b**

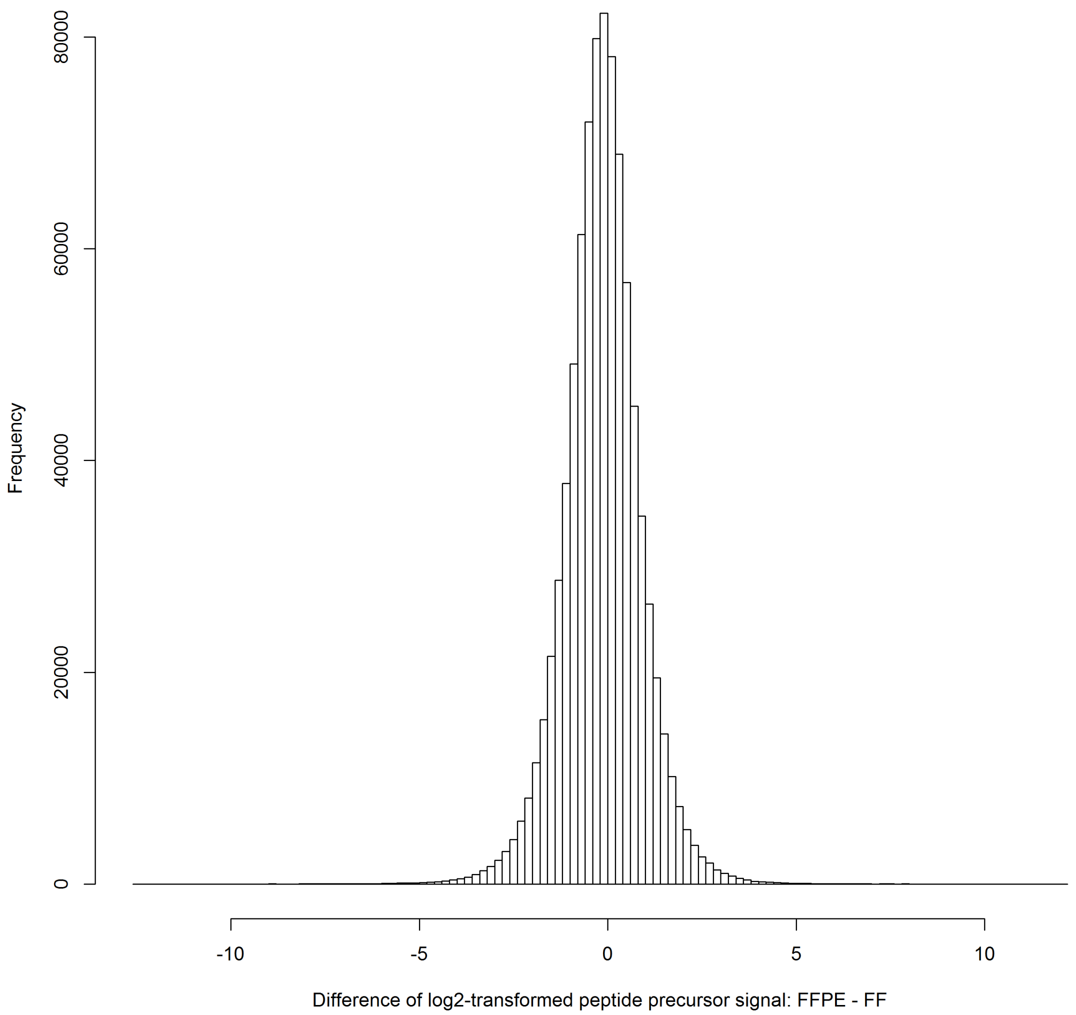


**Supplementary Figure 5b. Difference of log2-transformed peptide precursor signal in FF and FFPE samples in the PCF dataset.** This histogram shows the frequency of the value $log_2(FFPE) – log_2(FF)$ for each peptide precursor. Y-axis counts the number of peptide precursor in a tissue sample. It is clear that the distribution of the signal difference centered around 0, and exhibited two mirrored tails, indicating there is little systematic bias in the peptide quantities in FFPE or FF samples. In this plot, mean value is -0.1386(corresponding to a FFPE-to-FF ratio of 0.9084), median value is -0.1339 (corresponding to a FFPE-to-FF ratio of 0.9114). The 25% quantile is -0.7205 (corresponding to a FFPE-to-FF ratio of 0.6069), whereas the 75% quantile is 0.4507(corresponding to a FFPE-to-FF ratio of 1.3667). The data indicated that more than 50% of the peptide precursors showed a fold-change difference in their signal between FF and FFPE that was below 40%.

**Supplementary Figure 5c**

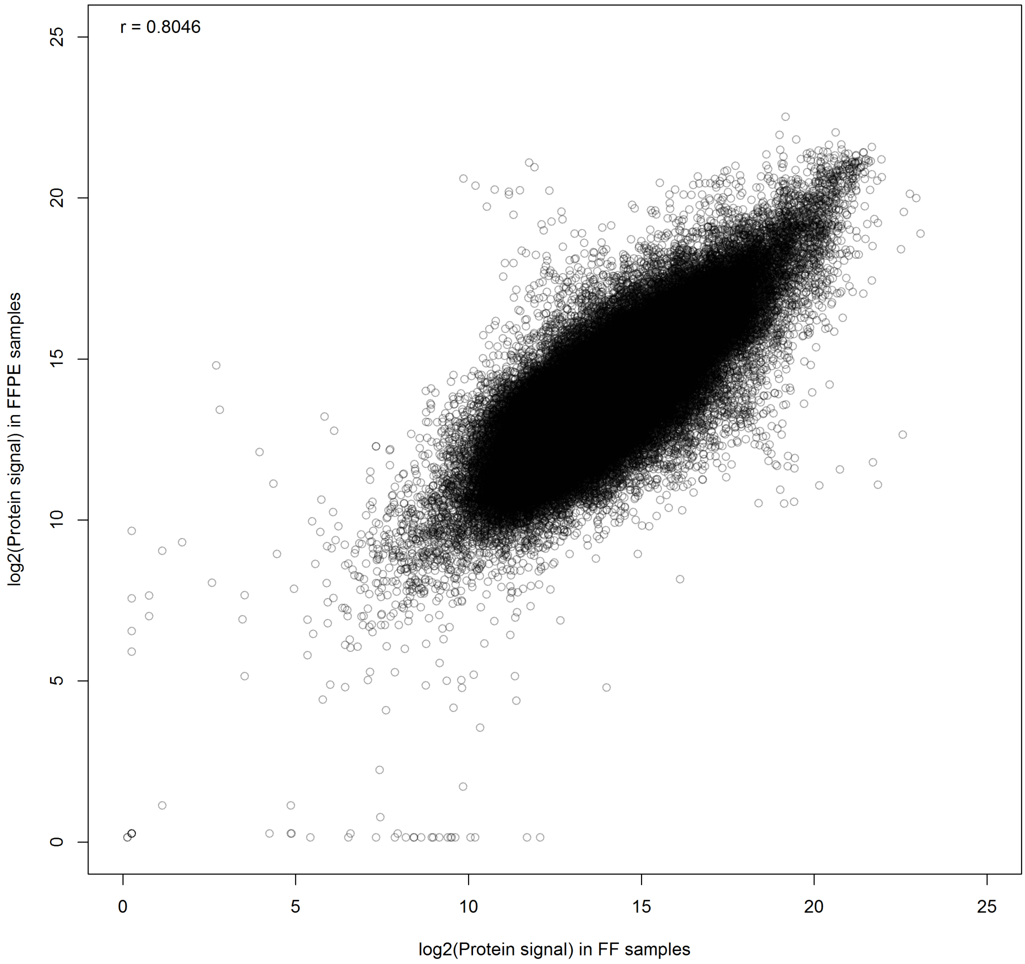


**Supplementary Figure 5c. Comparison of protein signal in FF and FFPE samples in the PCF dataset.** Each dot indicates a protein that was quantified in both FF (x-axis) and FFPE (y-axis) samples. Its log2-transformed signals are mapped in the two axes. Pearson correlation r=0.8046. The correlation at protein level is higher than that at peptide level, probably because we selected the best flyer peptide precursor for each protein thus reducing technical variation from peptides of low quantitative quality.

**Supplementary Figure 5d**

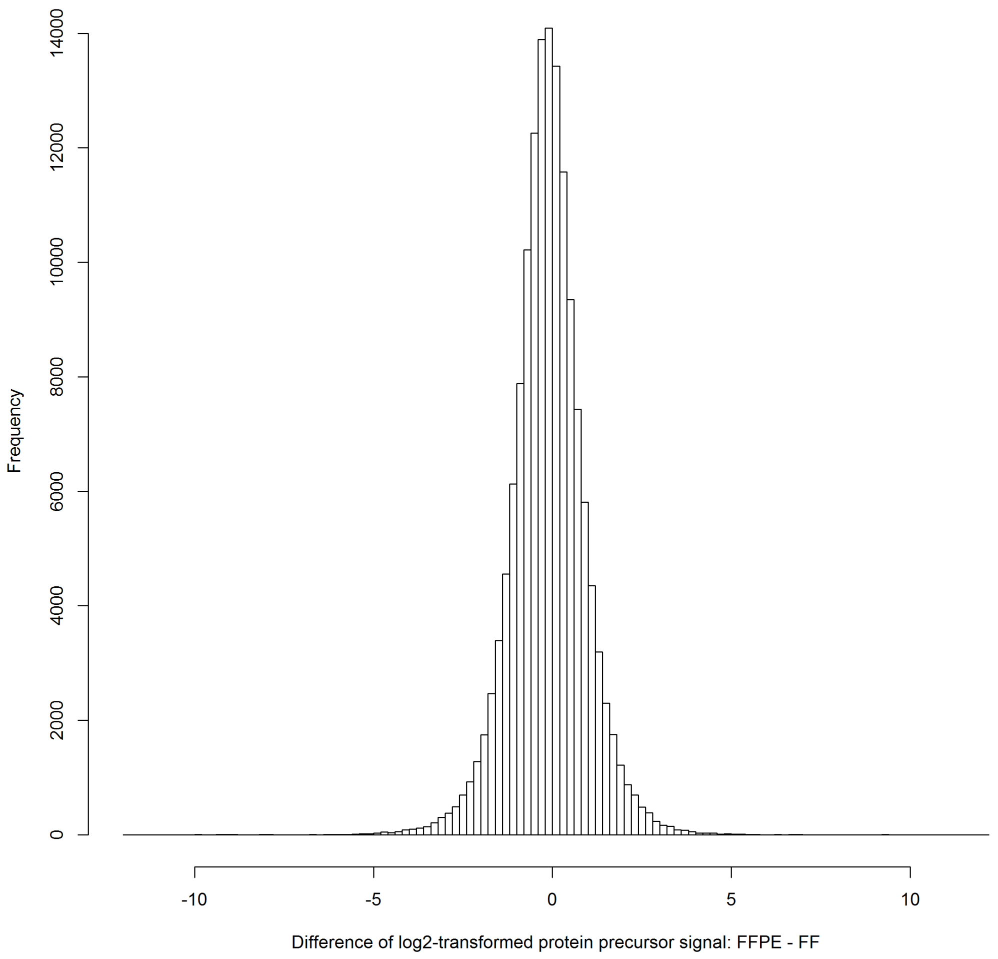


**Supplementary Figure 5d. Difference of log2-transformed protein signal in FF and FFPE samples in the PCF dataset.** This histogram shows the frequency of the value $\log_2(FFPE) - \log_2(FF)$ for each protein. Y-axis counts the number of protein in a tissue sample. The distribution appeared similar to peptide level, indicating the observed difference between FFPE and FF samples was consistent. The mean value is -0.1260 (corresponding to a FFPE-to-FF ratio of 0.9169), the median value is -0.1339 (corresponding to a FFPE-to-FF ratio of 0.9164). The 25% quantile is -0.6927 (corresponding to a FFPE-to-FF ratio of 0.6187), whereas the 75% quantile is 0.4495 (corresponding to a FFPE-to-FF ratio of 1.3656). The data indicate that 50% protein between FF and FFPE samples were below 39% difference.

#### Supplementary Figure 6

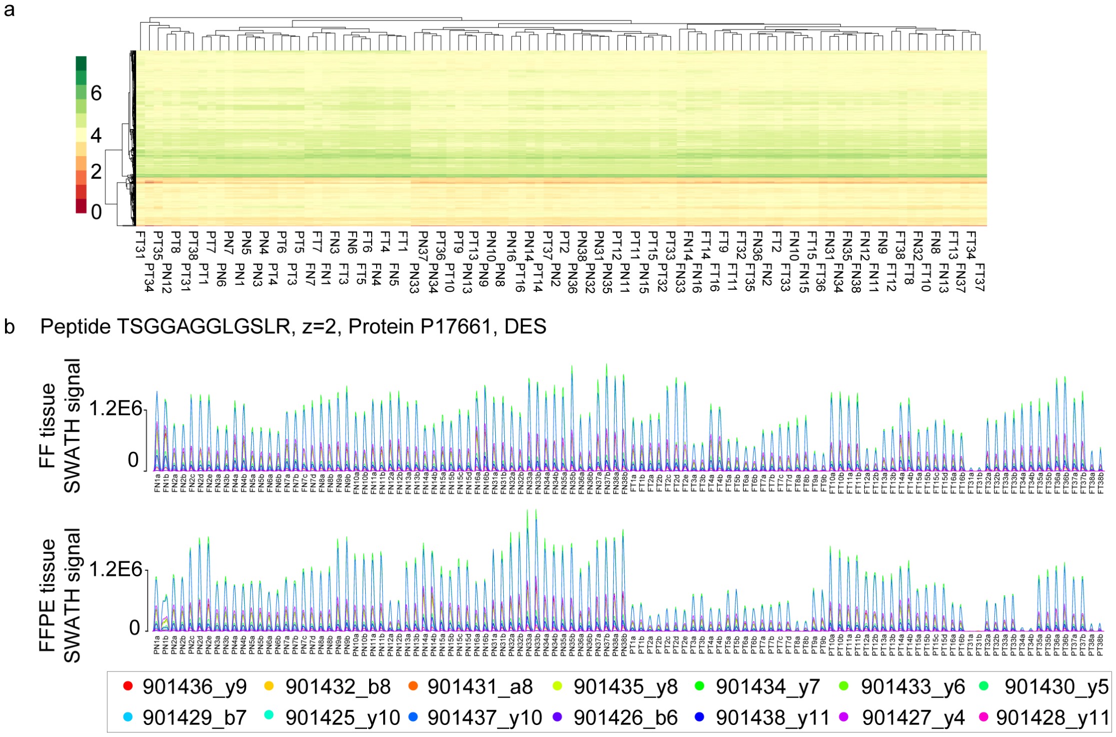


**Supplementary Figure 6. Comparison of fresh frozen and FFPE tissues in the PCF dataset.** (**a**) Cluster graph of proteins quantified in FFPE and FF tissues. Each row depicts the expression pattern of a protein for each sample tested. Each column represents a tissue sample. (**b**) Raw signals from DIA-expert for peptide TSGGAGGLGSLR (z=2) in protein DES. Upper panel: 110 FF samples; lower panel: 110 counterpart FFPE samples.

**Supplementary Figure 7**. **Effect of sample storage period on the proteome maps acquired by PCT-SWATH for both FF and FFPE samples.** The samples of the 24 ProCOC prostate cancer PCF dataset were selected to represent different periods of storage in either FF or FFPE state as detailed in **Supplementary Table 1**. 16 samples were procured before the year 2010 (referred to as "old" samples in this document), whereas 8 samples were procured after 2010 (referred to as "new" samples). The mean storage period for the old samples was 7.1 years, while the mean storage period for the new samples was 4.6 years (**a**).

We first checked if the TIC for each of the 224 SWATH runs at both MS1 (**b**) and MS2 (**c**) level. Surprisingly, we found the TIC at MS1 level segregated into two clear clusters. 80 SWATH files from the old samples exhibited substantially low MS1 compared to the other old and new samples (**h**). We found that these 80 SWATH files were all from the first batch of SWATH runs from 29 March 2016 to 1 April 2016, indicating a significant batch effect at MS1 level. Interestingly, this batch effect was not observed at MS2 level (**c**). We then computed the averaged MS2 TIC for FF (**d**) and FFPE (**e**) samples and confirmed that MS2 level results were not affected. All our downstream analysis was normalized and processed based on MS2 signals, therefore, we did not explore the observed MS1 instability issue any further. It was probably caused by instrument parameter deviation after cleaning and tuning.

In (**d)** and (**e**), we observed slightly higher MS2 TIC in old samples, however the TIC shapes were similar and such discrepancy is expected to be normalized in the downstream analysis. Therefore, we conclude that the storage time did not alter MS2 TIC.

We next compared the peptide precursors quantified in new and old FFPE and FF samples. As shown in **(f)**, the median signal in new samples are highly correlated with that in old samples (r=0.9735), indicating the sample storage time did not substantially alter peptide expression.

Median signal in new and old samples at protein level was analyzed in **g**. We observed an even higher Pearson correlation (r=0.9815).

In conclusion, we did not observe substantial difference between old and new samples at raw signal, peptide and protein level. However, it is worth noting that we did not compare tissue samples stored for over 10 years, which are not rare in biobanks, due to the lack of counterpart fresh frozen tissues stored for the same duration. Future research is required to more comprehensively investigate the influence of storage time.

**Supplementary Figure 7a**

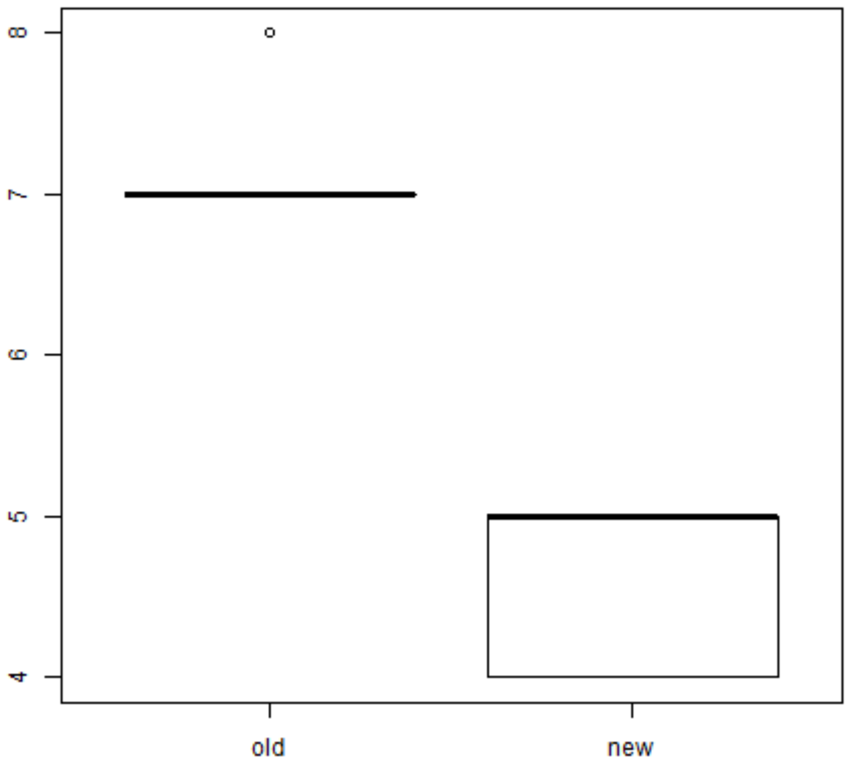


**Supplementary Figure 7a. Comparison of two sample groups with different storage periods.** Tissue samples from 16 patients were collected before the year 2010 (referred to as "old"), while samples from 8 patients were collected after the year 2010 (referred to as "new"). Welch Two Sample t-test using t.test function in R reported t = 12.381, df = 10.156, *p*-value = 1.871e-07. Mean storage period of old tissue iwas 7.1 years, while mean storage period of new tissue was 4.6 years.

#### Supplementary Figure 7b

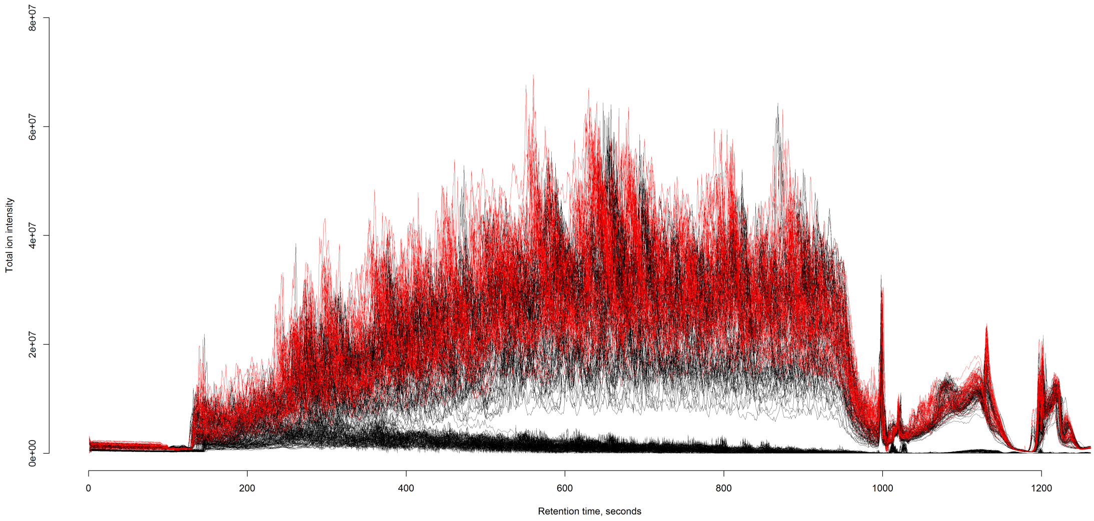


**Supplementary Figure 7b. MS1 TIC of 224 prostate tissue samples.** Red curves indicate new samples (collected after year 2010); while black curves indicate old samples (collected before year 2010).

#### Supplementary Figure 7c

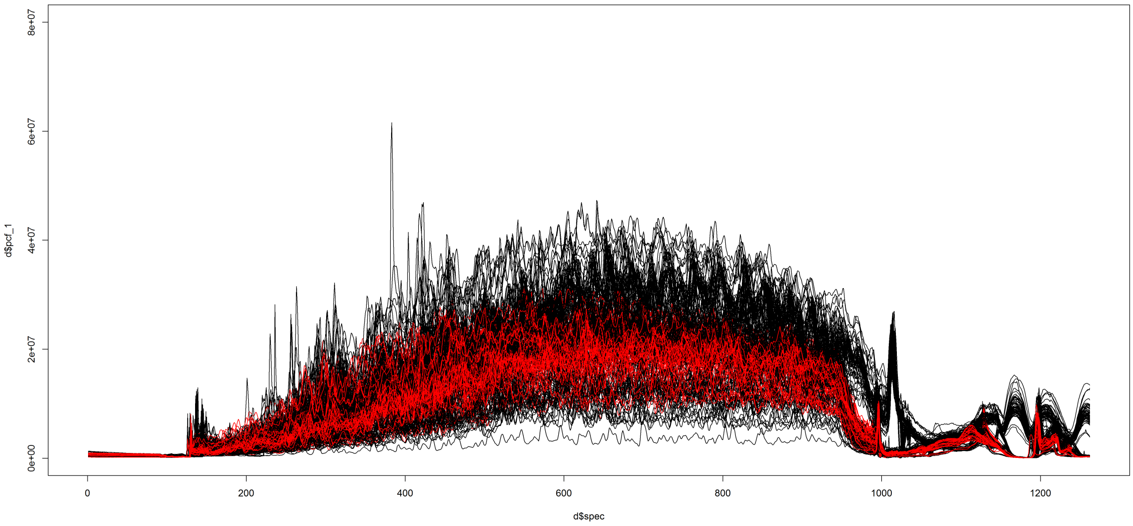


**Supplementary Figure 7c. MS2 TIC of 224 prostate tissue samples.** Red curves indicate new samples (collected after year 2010); while black curves indicate old samples (collected before year 2010).

#### Supplementary Figure 7d

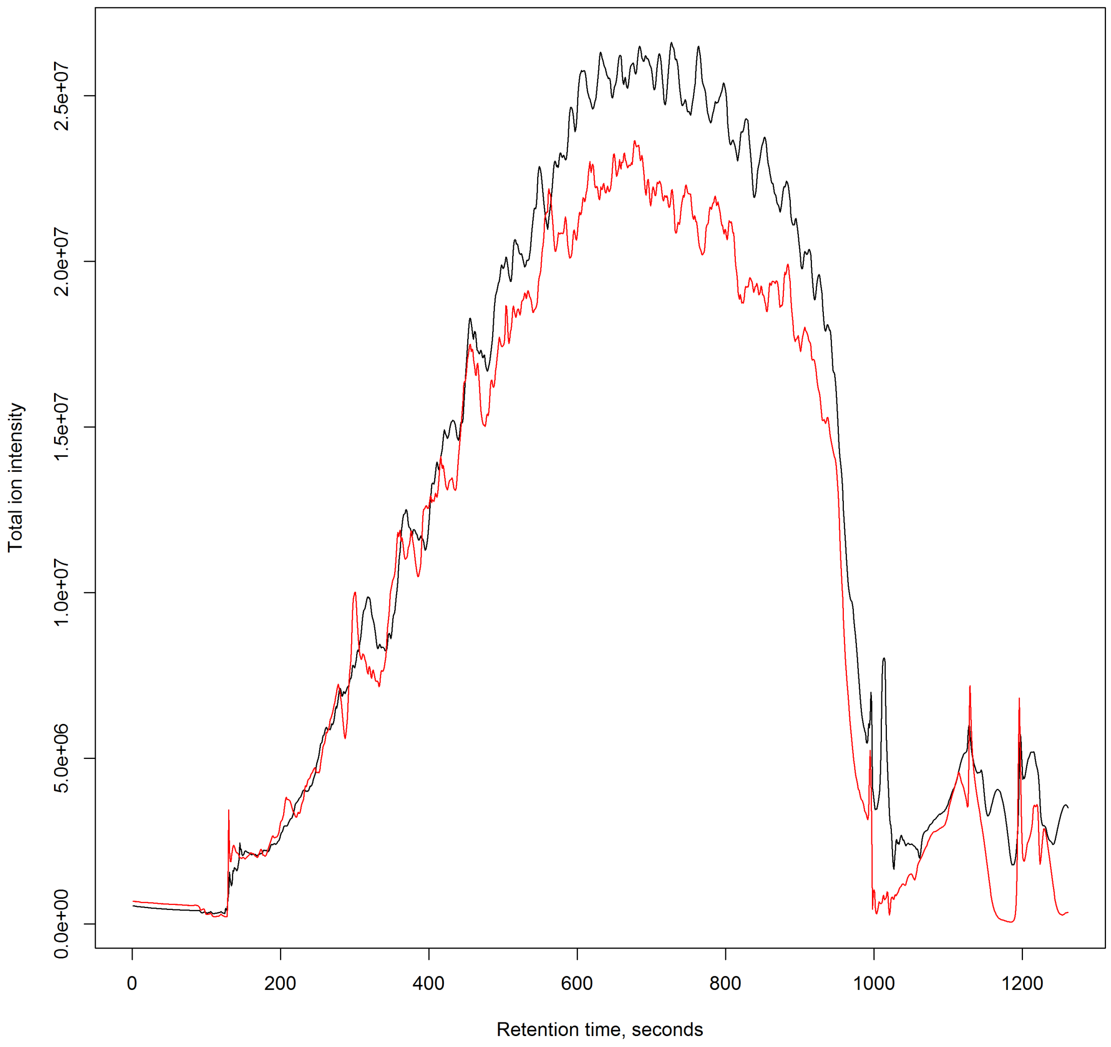


**Supplementary Figure 7d. Comparison of averaged MS2 TIC of old (black curve) and new (red curve) FF samples.** x-axis indicates the liquid chromatogram retention time in seconds. y-axis indicates the averaged MS2 total ion intensity from all new or old samples.

**Supplementary Figure 7e**

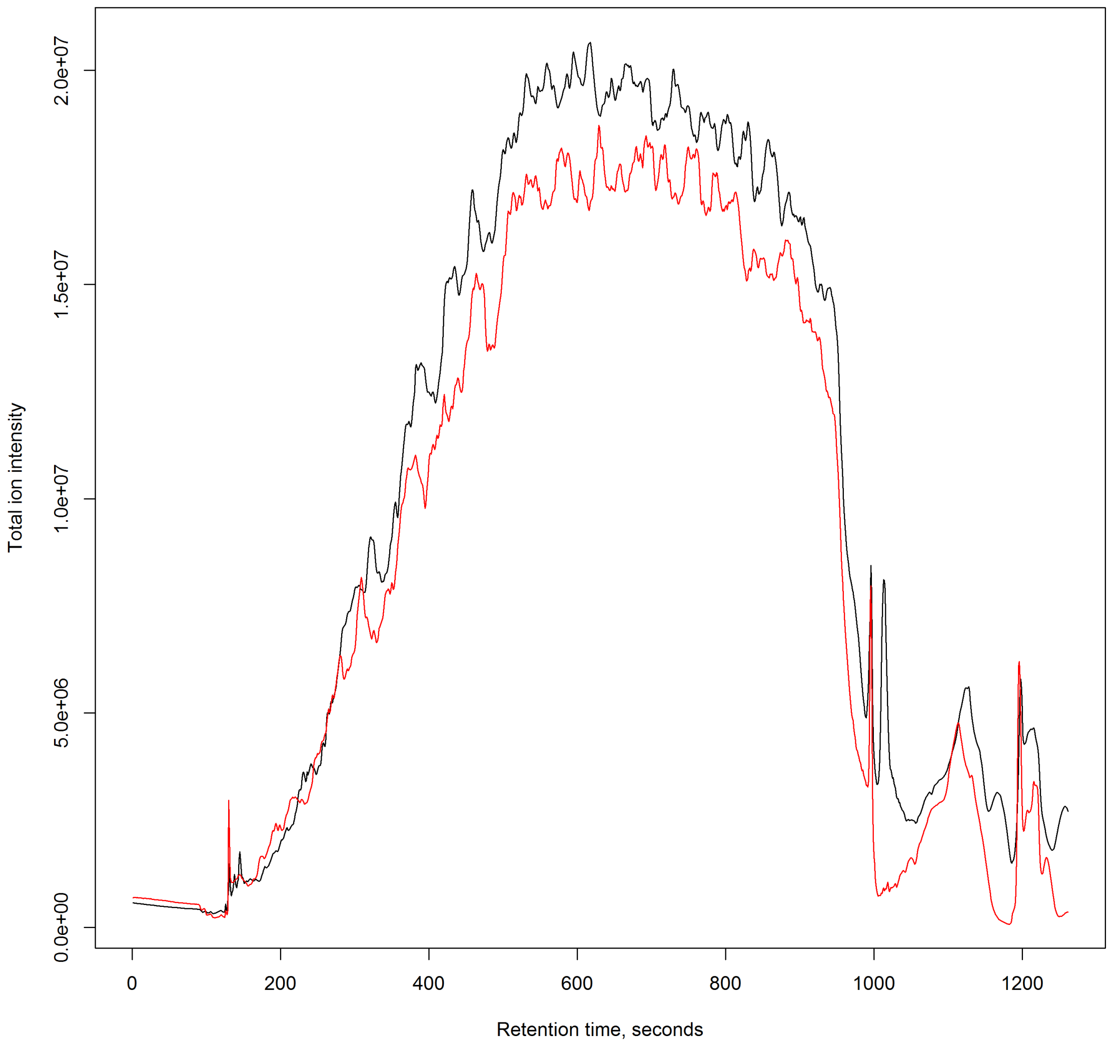


**Supplementary Figure 7e. Comparison of averaged MS2 TIC of old (black curve) and new (red curve) FFPE samples.** x-axis indicates the liquid chromatogram retention time in seconds. y-axis indicates the averaged MS2 total ion intensity from all new or old samples.

**Supplementary Figure 7f**

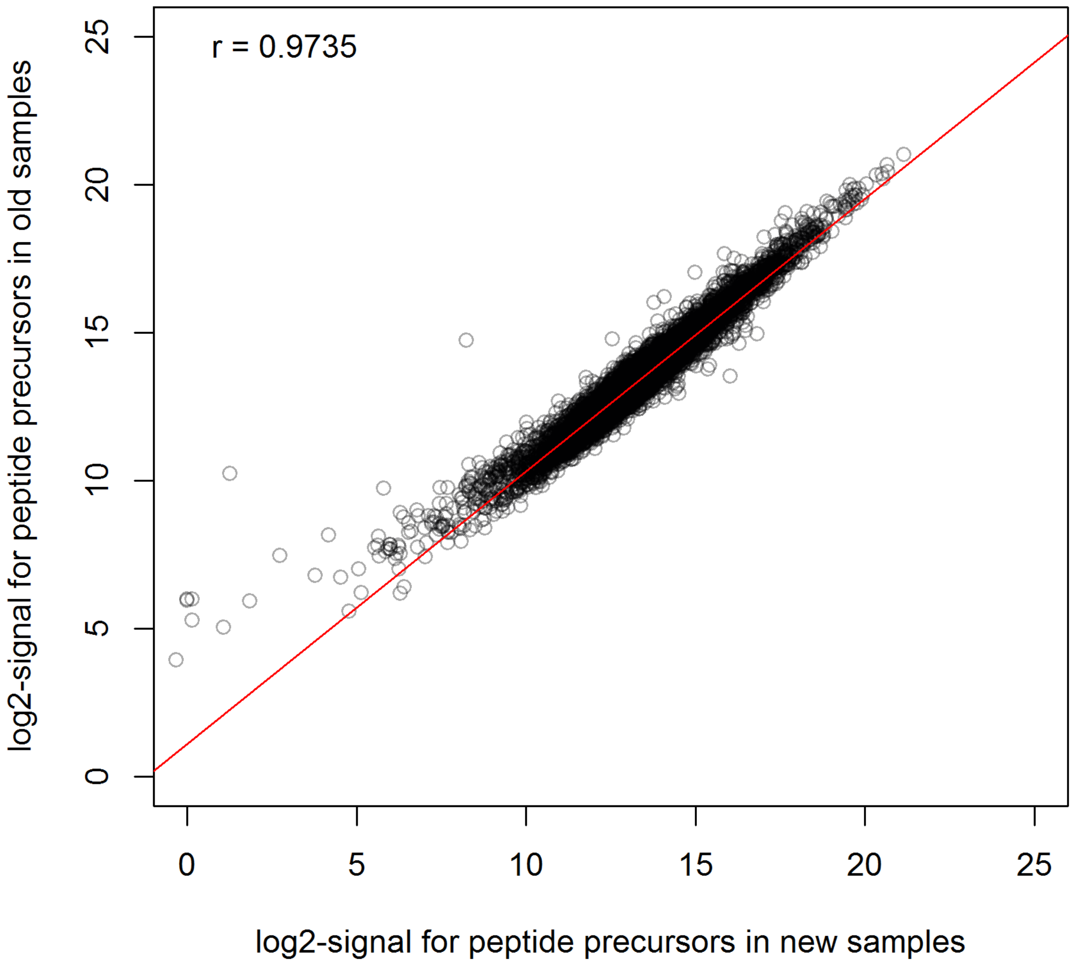


**Supplementary Figure 7f. Comparison of new and old samples at peptide level.** x-axis indicates the median log2 transformed peptide precursor signal in new samples (collected after year 2010); y-axis indicates the median log2 transformed peptide precursor signal in old samples (collected before year 2010). A total of 18,129 peptide precursors are shown. Pearson correlation coefficient r=0.9735.

**Supplementary Figure 7g**

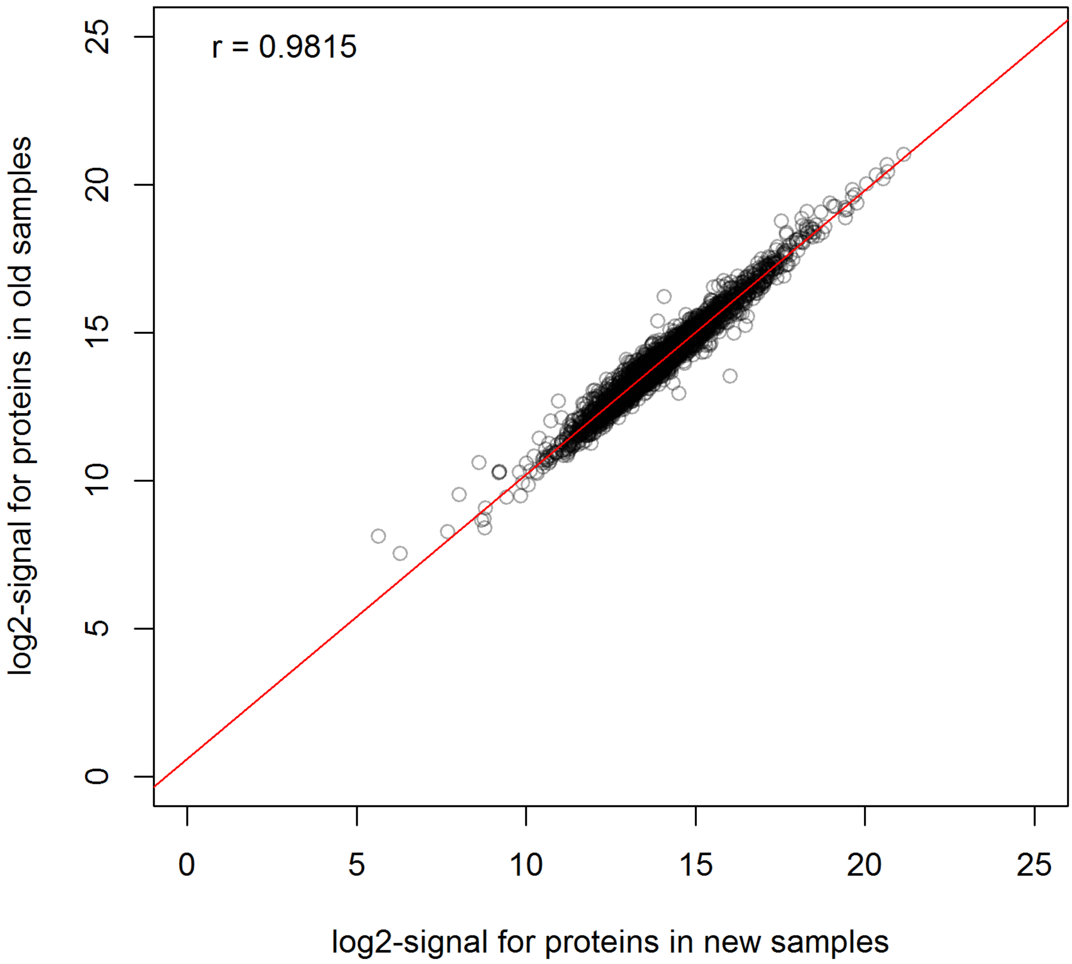


**Supplementary Figure 7g. Comparison of new and old samples at protein level.** x-axis indicates the median log2 transformed protein signal in new samples (collected after year 2010); y-axis indicates the median log2 transformed protein signal in old samples (collected before year 2010). A total of 3,030 proteins are shown. Pearson correlation coefficient r=0.9815.

#### Supplementary Figure 7h.

| pcf_id | sample_age | MS1_TIC | MS2_TIC | sample_name | storage_type | normal_tumor | patient_id | sample_name2 | tech_rep | MS_batch |
|---|---|---|---|---|---|---|---|---|---|---|
| pcf183 | old | 587384550 | 13758321092 | PT6b | P | T | 6 | PT6 | b | L160329 |
| pcf182 | old | 689579820 | 14038549192 | PT6a | P | T | 6 | PT6 | a | L160329 |
| pcf17 | old | 751640115 | 19391016430 | FN7b | F | N | 7 | FN7 | b | L160329 |
| pcf207 | old | 774450629 | 15596561865 | PT16b | P | T | 16 | PT16 | b | L160401 |
| pcf203 | old | 782951452 | 15529535580 | PT15b | P | T | 15 | PT15 | b | L160401 |
| pcf181 | old | 817724223 | 9625013654 | PT5b | P | T | 5 | PT5 | b | L160329 |
| pcf180 | old | 846725733 | 9584650123 | PT5a | P | T | 5 | PT5 | a | L160329 |
| pcf16 | old | 854406744 | 19339060694 | FN7a | F | N | 7 | FN7 | a | L160329 |
| pcf206 | old | 870237402 | 15893091711 | PT16a | P | T | 16 | PT16 | a | L160401 |
| pcf202 | old | 903919569 | 15587530646 | PT15a | P | T | 15 | PT15 | a | L160401 |
| pcf72 | old | 908857019 | 20224327196 | FT7b | F | T | 7 | FT7 | b | L160329 |
| pcf179 | old | 916516043 | 10408975162 | PT4b | P | T | 4 | PT4 | b | L160329 |
| pcf185 | old | 955511235 | 25861984597 | PT7b | P | T | 7 | PT7 | b | L160329 |
| pcf38 | old | 962256486 | 16039048421 | FN16a | F | N | 16 | FN16 | a | L160401 |
| pcf39 | old | 995359510 | 16689754597 | FN16b | F | N | 16 | FN16 | b | L160401 |
| pcf94 | old | 1004669622 | 13989852390 | FT16b | F | T | 16 | FT16 | b | L160401 |
| pcf178 | old | 1007214180 | 10445396383 | PT4a | P | T | 4 | PT4 | a | L160329 |
| pcf15 | old | 1030435598 | 19222057458 | FN6b | F | N | 6 | FN6 | b | L160329 |
| pcf90 | old | 1050632797 | 15190463524 | FT15b | F | T | 15 | FT15 | b | L160401 |
| pcf93 | old | 1055798449 | 13889369927 | FT16a | F | T | 16 | FT16 | a | L160401 |
| pcf70 | old | 1056614526 | 17597886278 | FT6b | F | T | 6 | FT6 | b | L160329 |
| pcf184 | old | 1062342119 | 25384298064 | PT7a | P | T | 7 | PT7 | a | L160329 |
| pcf71 | old | 1072123222 | 20188655764 | FT7a | F | T | 7 | FT7 | a | L160329 |
| pcf35 | old | 1077240013 | 17706618661 | FN15b | F | N | 15 | FN15 | b | L160401 |
| pcf150 | old | 1085871983 | 14537775118 | PN16b | P | N | 16 | PN16 | b | L160401 |
| pcf176 | old | 1111840352 | 13257366445 | PT3a | P | T | 3 | PT3 | a | L160329 |
| pcf34 | old | 1112742767 | 16734715226 | FN15a | F | N | 15 | FN15 | a | L160401 |
| pcf128 | old | 1119414131 | 16322574829 | PN7b | P | N | 7 | PN7 | b | L160329 |
| pcf89 | old | 1124391842 | 14845994551 | FT15a | F | T | 15 | FT15 | a | L160401 |
| pcf201 | old | 1124404688 | 13035477108 | PT14b | P | T | 14 | PT14 | b | L160401 |
| pcf14 | old | 1132781101 | 19465789294 | FN6a | F | N | 6 | FN6 | a | L160329 |
| pcf69 | old | 1152654409 | 17767341224 | FT6a | F | T | 6 | FT6 | a | L160329 |
| pcf177 | old | 1187580335 | 13144607669 | PT3b | P | T | 3 | PT3 | b | L160329 |
| pcf33 | old | 1200849257 | 15127552101 | FN14b | F | N | 14 | FN14 | b | L160401 |
| pcf149 | old | 1210213392 | 15460106189 | PN16a | P | N | 16 | PN16 | a | L160401 |
| pcf88 | old | 1264693326 | 16644720946 | FT14b | F | T | 14 | FT14 | b | L160401 |
| pcf146 | old | 1281282801 | 15946054008 | PN15b | P | N | 15 | PN15 | b | L160401 |
| pcf127 | old | 1287685670 | 15851257583 | PN7a | P | N | 7 | PN7 | a | L160329 |
| pcf200 | old | 1344389355 | 13267453461 | PT14a | P | T | 14 | PT14 | a | L160401 |
| pcf87 | old | 1351527840 | 16558461871 | FT14a | F | T | 14 | FT14 | a | L160401 |
| pcf145 | old | 1369863887 | 14638823124 | PN15a | P | N | 15 | PN15 | a | L160401 |
| pcf170 | old | 1395653897 | 18485798326 | PT2b | P | T | 2 | PT2 | b | L160329 |
| pcf144 | old | 1417768267 | 13354446922 | PN14b | P | N | 14 | PN14 | b | L160401 |
| pcf169 | old | 1428542409 | 18987699475 | PT2a | P | T | 2 | PT2 | a | L160329 |
| pcf13 | old | 1473942887 | 19214951510 | FN5b | F | N | 5 | FN5 | b | L160329 |
| pcf32 | old | 1494927657 | 15948939687 | FN14a | F | N | 14 | FN14 | a | L160401 |
| pcf12 | old | 1523633160 | 19128137675 | FN5a | F | N | 5 | FN5 | a | L160329 |
| pcf124 | old | 1525425364 | 17371101168 | PN5b | P | N | 5 | PN5 | b | L160329 |
| pcf68 | old | 1546787443 | 18908706051 | FT5b | F | T | 5 | FT5 | b | L160329 |
| pcf67 | old | 1553928012 | 18536768151 | FT5a | F | T | 5 | FT5 | a | L160329 |
| pcf11 | old | 1557374892 | 17465396130 | FN4b | F | N | 4 | FN4 | b | L160329 |
| pcf126 | old | 1564276176 | 25368131927 | PN6b | P | N | 6 | PN6 | b | L160329 |
| pcf123 | old | 1569670301 | 17366752745 | PN5a | P | N | 5 | PN5 | a | L160329 |
| pcf168 | old | 1590372102 | 22187520563 | PT1b | P | T | 1 | PT1 | b | L160329 |
| pcf4 | old | 1601506192 | 19267557180 | FN2b | F | N | 2 | FN2 | b | L160329 |
| pcf2 | old | 1629758744 | 24412326938 | FN1b | F | N | 1 | FN1 | b | L160329 |
| pcf3 | old | 1631651761 | 19465662383 | FN2a | F | N | 2 | FN2 | a | L160329 |
| pcf66 | old | 1660948374 | 19644052905 | FT4b | F | T | 4 | FT4 | b | L160329 |
| pcf10 | old | 1676968290 | 17392328814 | FN4a | F | N | 4 | FN4 | a | L160329 |
| pcf64 | old | 1705412276 | 18634928886 | FT3b | F | T | 3 | FT3 | b | L160329 |
| pcf65 | old | 1708265966 | 19669423110 | FT4a | F | T | 4 | FT4 | a | L160329 |
| pcf143 | old | 1710038456 | 13508077838 | PN14a | P | N | 14 | PN14 | a | L160401 |
| pcf122 | old | 1716703966 | 18173711893 | PN4b | P | N | 4 | PN4 | b | L160329 |
| pcf63 | old | 1716873682 | 18601843352 | FT3a | F | T | 3 | FT3 | a | L160329 |
| pcf57 | old | 1725088845 | 21273875627 | FT1b | F | T | 1 | FT1 | b | L160329 |
| pcf8 | old | 1740214151 | 19791170295 | FN3a | F | N | 3 | FN3 | a | L160329 |
| pcf125 | old | 1786406978 | 25383221327 | PN6a | P | N | 6 | PN6 | a | L160329 |
| pcf9 | old | 1799135521 | 19727750996 | FN3b | F | N | 3 | FN3 | b | L160329 |
| pcf1 | old | 1831592143 | 24133929018 | FN1a | F | N | 1 | FN1 | a | L160329 |
| pcf121 | old | 1833147788 | 17695884549 | PN4a | P | N | 4 | PN4 | a | L160329 |
| pcf113 | old | 1847613885 | 15606217516 | PN2a | P | N | 2 | PN2 | a | L160329 |
| pcf59 | old | 1853118816 | 18374238187 | FT2b | F | T | 2 | FT2 | b | L160329 |
| pcf119 | old | 1871471744 | 16915706857 | PN3a | P | N | 3 | PN3 | a | L160329 |
| pcf56 | old | 1898682449 | 21677403143 | FT1a | F | T | 1 | FT1 | a | L160329 |
| pcf167 | old | 1903011136 | 22434636486 | PT1a | P | T | 1 | PT1 | a | L160329 |
| pcf114 | old | 1905082032 | 15401203291 | PN2b | P | N | 2 | PN2 | b | L160329 |
| pcf58 | old | 1920528976 | 18564557410 | FT2a | F | T | 2 | FT2 | a | L160329 |
| pcf112 | old | 1963723510 | 18430814644 | PN1b | P | N | 1 | PN1 | b | L160329 |
| pcf120 | old | 2004985307 | 16059729162 | PN3b | P | N | 3 | PN3 | b | L160329 |
| pcf111 | old | 3388291604 | 17802541892 | PN1a | P | N | 1 | PN1 | a | L160329 |
| pcf118 | old | 7139256675 | 2494877656 | PN2f | P | N | 2 | PN2 | f | L160606 |
| pcf134 | old | 7826101086 | 5118640178 | PN9b | P | N | 9 | PN9 | b | L160526 |
| pcf133 | old | 8008775040 | 5323681687 | PN9a | P | N | 9 | PN9 | a | L160526 |
| pcf193 | old | 8508744890 | 5503561591 | PT10b | P | T | 10 | PT10 | b | L160526 |
| pcf192 | old | 8994383585 | 5957205223 | PT10a | P | T | 10 | PT10 | a | L160526 |
| pcf131 | old | 1.0075E+10 | 7164642163 | PN8a | P | N | 8 | PN8 | a | L160526 |
| pcf136 | old | 1.026E+10 | 6778359802 | PN10b | P | N | 10 | PN10 | b | L160526 |
| pcf132 | old | 1.0556E+10 | 7429378535 | PN8b | P | N | 8 | PN8 | b | L160526 |
| pcf135 | old | 1.0979E+10 | 7412086536 | PN10a | P | N | 10 | PN10 | a | L160526 |
| pcf188 | old | 1.1139E+10 | 7724200127 | PT8a | P | T | 8 | PT8 | a | L160526 |
| pcf175 | old | 1.1215E+10 | 4825678020 | PT2g | P | T | 2 | PT2 | g | L160606 |

| | | | | | | | | | | |
|---|---|---|---|---|---|---|---|---|---|---|
| **pcf138** | old | 1.1508E+10 | 7563904484 | PN11b | P | N | 11 | PN11 | b | L160526 |
| **pcf137** | old | 1.1664E+10 | 7720900674 | PN11a | P | N | 11 | PN11 | a | L160526 |
| **pcf189** | old | 1.1791E+10 | 8047142794 | PT8b | P | T | 8 | PT8 | b | L160526 |
| **pcf162** | new | 1.1803E+10 | 6454892252 | PN36b | P | N | 36 | PN36 | b | L160530 |
| **pcf117** | old | 1.1833E+10 | 4983893007 | PN2e | P | N | 2 | PN2 | e | L160606 |
| **pcf140** | old | 1.2017E+10 | 7781542801 | PN12b | P | N | 12 | PN12 | b | L160526 |
| **pcf23** | old | 1.2053E+10 | 7464307291 | FN9b | F | N | 9 | FN9 | b | L160526 |
| **pcf161** | new | 1.2073E+10 | 7172220861 | PN36a | P | N | 36 | PN36 | a | L160530 |
| **pcf195** | old | 1.2105E+10 | 7933550645 | PT11b | P | T | 11 | PT11 | b | L160526 |
| **pcf83** | old | 1.2199E+10 | 7419647840 | FT12a | F | T | 12 | FT12 | a | L160526 |
| **pcf139** | old | 1.2321E+10 | 8044944205 | PN12a | P | N | 12 | PN12 | a | L160526 |
| **pcf174** | old | 1.2631E+10 | 5683119597 | PT2f | P | T | 2 | PT2 | f | L160606 |
| **pcf84** | old | 1.2723E+10 | 7657192613 | FT12b | F | T | 12 | FT12 | b | L160526 |
| **pcf129** | old | 1.2751E+10 | 9470910566 | PN7c | P | N | 7 | PN7 | c | L160526 |
| **pcf173** | old | 1.3052E+10 | 5957777702 | PT2e | P | T | 2 | PT2 | e | L160606 |
| **pcf172** | old | 1.3376E+10 | 7197317341 | PT2d | P | T | 2 | PT2 | d | L160530 |
| **pcf116** | old | 1.3417E+10 | 6804980462 | PN2d | P | N | 2 | PN2 | d | L160530 |
| **pcf194** | old | 1.3481E+10 | 9049431024 | PT11a | P | T | 11 | PT11 | a | L160526 |
| **pcf115** | old | 1.3754E+10 | 7566939003 | PN2c | P | N | 2 | PN2 | c | L160530 |
| **pcf130** | old | 1.4229E+10 | 10461177463 | PN7d | P | N | 7 | PN7 | d | L160526 |
| **pcf154** | new | 1.4485E+10 | 8689531422 | PN32b | P | N | 32 | PN32 | b | L160529 |
| **pcf153** | new | 1.4596E+10 | 8986702964 | PN32a | P | N | 32 | PN32 | a | L160529 |
| **pcf197** | old | 1.4872E+10 | 9486243042 | PT12b | P | T | 12 | PT12 | b | L160526 |
| **pcf171** | old | 1.4938E+10 | 8826665114 | PT2c | P | T | 2 | PT2 | c | L160530 |
| **pcf215** | new | 1.515E+10 | 7191464779 | PT34b | P | T | 34 | PT34 | b | L160606 |
| **pcf196** | old | 1.5473E+10 | 9906134533 | PT12a | P | T | 12 | PT12 | a | L160526 |
| **pcf222** | new | 1.5627E+10 | 8766124007 | PT37b | P | T | 37 | PT37 | b | L160530 |
| **pcf216** | new | 1.59E+10 | 7513765123 | PT34c | P | T | 34 | PT34 | c | L160606 |
| **pcf221** | new | 1.5979E+10 | 9597261467 | PT37a | P | T | 37 | PT37 | a | L160530 |
| **pcf160** | new | 1.6116E+10 | 9256384773 | PN35b | P | N | 35 | PN35 | b | L160530 |
| **pcf151** | new | 1.6182E+10 | 10303003166 | PN31a | P | N | 31 | PN31 | a | L160529 |
| **pcf152** | new | 1.6458E+10 | 10474446503 | PN31b | P | N | 31 | PN31 | b | L160529 |
| **pcf61** | old | 1.6722E+10 | 6797973170 | FT2d | F | T | 2 | FT2 | d | L160606 |
| **pcf164** | new | 1.6898E+10 | 9401162921 | PN37b | P | N | 37 | PN37 | b | L160530 |
| **pcf163** | new | 1.7108E+10 | 10180645486 | PN37a | P | N | 37 | PN37 | a | L160530 |
| **pcf214** | new | 1.7266E+10 | 10936068432 | PT34a | P | T | 34 | PT34 | a | L160530 |
| **pcf159** | new | 1.7339E+10 | 10673643192 | PN35a | P | N | 35 | PN35 | a | L160530 |
| **pcf6** | old | 1.7584E+10 | 7449363169 | FN2d | F | N | 2 | FN2 | d | L160606 |
| **pcf7** | old | 1.7652E+10 | 7423576047 | FN2e | F | N | 2 | FN2 | e | L160606 |
| **pcf62** | old | 1.7808E+10 | 7266270838 | FT2e | F | T | 2 | FT2 | e | L160606 |
| **pcf49** | new | 1.7819E+10 | 8891038782 | FN35b | F | N | 35 | FN35 | b | L160530 |
| **pcf198** | old | 1.7838E+10 | 11869562201 | PT13a | P | T | 13 | PT13 | a | L160526 |
| **pcf212** | new | 1.8102E+10 | 11437859747 | PT33a | P | T | 33 | PT33 | a | L160529 |
| **pcf213** | new | 1.8111E+10 | 11408531117 | PT33b | P | T | 33 | PT33 | b | L160529 |
| **pcf166** | new | 1.8425E+10 | 10225340033 | PN38b | P | N | 38 | PN38 | b | L160530 |
| **pcf209** | new | 1.846E+10 | 11977920407 | PT31b | P | T | 31 | PT31 | b | L160529 |
| **pcf208** | new | 1.8527E+10 | 12044606934 | PT31a | P | T | 31 | PT31 | a | L160529 |
| **pcf156** | new | 1.8583E+10 | 11252405419 | PN33b | P | N | 33 | PN33 | b | L160529 |
| **pcf60** | old | 1.8801E+10 | 9602959015 | FT2c | F | T | 2 | FT2 | c | L160530 |
| **pcf199** | old | 1.8845E+10 | 12456226175 | PT13b | P | T | 13 | PT13 | b | L160526 |
| **pcf142** | old | 1.8985E+10 | 12393372522 | PN13b | P | N | 13 | PN13 | b | L160526 |
| **pcf53** | new | 1.9011E+10 | 9241320488 | FN37b | F | N | 37 | FN37 | b | L160530 |
| **pcf26** | old | 1.9022E+10 | 11311384201 | FN11a | F | N | 11 | FN11 | a | L160526 |
| **pcf155** | new | 1.9101E+10 | 11532962729 | PN33a | P | N | 33 | PN33 | a | L160529 |
| **pcf22** | old | 1.9198E+10 | 11755449402 | FN9a | F | N | 9 | FN9 | a | L160526 |
| **pcf30** | old | 1.9441E+10 | 11350229021 | FN13a | F | N | 13 | FN13 | a | L160526 |
| **pcf165** | new | 1.9529E+10 | 11543441425 | PN38a | P | N | 38 | PN38 | a | L160530 |
| **pcf5** | old | 1.9609E+10 | 10350400990 | FN2c | F | N | 2 | FN2 | c | L160530 |
| **pcf224** | new | 1.9766E+10 | 11105062228 | PT38b | P | T | 38 | PT38 | b | L160530 |
| **pcf148** | old | 1.9874E+10 | 12996608829 | PN15d | P | N | 15 | PN15 | d | L160526 |
| **pcf223** | new | 2.0106E+10 | 12023214179 | PT38a | P | T | 38 | PT38 | a | L160530 |
| **pcf147** | old | 2.0174E+10 | 13122646274 | PN15c | P | N | 15 | PN15 | c | L160526 |
| **pcf79** | old | 2.027E+10 | 12001083500 | FT10a | F | T | 10 | FT10 | a | L160526 |
| **pcf98** | new | 2.0494E+10 | 11688281263 | FT32b | F | T | 32 | FT32 | b | L160529 |
| **pcf31** | old | 2.0513E+10 | 11908421783 | FN13b | F | N | 13 | FN13 | b | L160526 |
| **pcf27** | old | 2.0585E+10 | 12079125525 | FN11b | F | N | 11 | FN11 | b | L160526 |
| **pcf190** | old | 2.059E+10 | 14430839298 | PT9a | P | T | 9 | PT9 | a | L160526 |
| **pcf19** | old | 2.062E+10 | 12369928803 | FN7d | F | N | 7 | FN7 | d | L160526 |
| **pcf29** | old | 2.0652E+10 | 12026983736 | FN12b | F | N | 12 | FN12 | b | L160526 |
| **pcf191** | old | 2.0739E+10 | 14431050060 | PT9b | P | T | 9 | PT9 | b | L160526 |
| **pcf85** | old | 2.0783E+10 | 12339577614 | FT13a | F | T | 13 | FT13 | a | L160526 |
| **pcf77** | old | 2.0895E+10 | 12257124075 | FT9a | F | T | 9 | FT9 | a | L160526 |
| **pcf18** | old | 2.0943E+10 | 12584155282 | FN7c | F | N | 7 | FN7 | c | L160526 |
| **pcf41** | new | 2.1076E+10 | 11884663933 | FN31b | F | N | 31 | FN31 | b | L160529 |
| **pcf40** | new | 2.1113E+10 | 11958138446 | FN31a | F | N | 31 | FN31 | a | L160529 |
| **pcf74** | old | 2.1133E+10 | 12946269087 | FT7d | F | T | 7 | FT7 | d | L160526 |
| **pcf28** | old | 2.1292E+10 | 12459273053 | FN12a | F | N | 12 | FN12 | a | L160526 |
| **pcf73** | old | 2.1428E+10 | 12981805786 | FT7c | F | T | 7 | FT7 | c | L160526 |
| **pcf218** | new | 2.1469E+10 | 12749486090 | PT35b | P | T | 35 | PT35 | b | L160530 |
| **pcf141** | old | 2.1493E+10 | 13807350194 | PN13a | P | N | 13 | PN13 | a | L160526 |
| **pcf46** | new | 2.1561E+10 | 11531690079 | FN34a | F | N | 34 | FN34 | a | L160530 |
| **pcf95** | new | 2.1621E+10 | 13221219823 | FT31a | F | T | 31 | FT31 | a | L160529 |
| **pcf220** | new | 2.1682E+10 | 12371095565 | PT36b | P | T | 36 | PT36 | b | L160530 |
| **pcf210** | new | 2.1786E+10 | 14048072793 | PT32a | P | T | 32 | PT32 | a | L160529 |
| **pcf96** | new | 2.1798E+10 | 13200675499 | FT31b | F | T | 31 | FT31 | b | L160529 |
| **pcf101** | new | 2.1844E+10 | 12002222991 | FT34a | F | T | 34 | FT34 | a | L160530 |
| **pcf25** | old | 2.185E+10 | 13414173995 | FN10b | F | N | 10 | FN10 | b | L160526 |
| **pcf102** | new | 2.1965E+10 | 11477859160 | FT34b | F | T | 34 | FT34 | b | L160530 |
| **pcf86** | old | 2.1966E+10 | 12851820862 | FT13b | F | T | 13 | FT13 | b | L160526 |
| **pcf204** | old | 2.2124E+10 | 14843034930 | PT15c | P | T | 15 | PT15 | c | L160526 |
| **pcf205** | old | 2.219E+10 | 14888079198 | PT15d | P | T | 15 | PT15 | d | L160526 |
| **pcf217** | new | 2.2207E+10 | 13906473668 | PT35a | P | T | 35 | PT35 | a | L160530 |
| **pcf211** | new | 2.2229E+10 | 14270654862 | PT32b | P | T | 32 | PT32 | b | L160529 |
| **pcf186** | old | 2.2284E+10 | 15816815994 | PT7c | P | T | 7 | PT7 | c | L160526 |
| **pcf51** | new | 2.2294E+10 | 11399382865 | FN36b | F | N | 36 | FN36 | b | L160530 |
| **pcf187** | old | 2.231E+10 | 15916379463 | PT7d | P | T | 7 | PT7 | d | L160526 |

| | | | | | | | | | | |
|---|---|---|---|---|---|---|---|---|---|---|
| **pcf78** | old | 2.2311E+10 | 12958083020 | FT9b | F | T | 9 | FT9 | b | L160526 |
| **pcf24** | old | 2.2548E+10 | 13924537831 | FN10a | F | N | 10 | FN10 | a | L160526 |
| **pcf75** | old | 2.2613E+10 | 13845264929 | FT8a | F | T | 8 | FT8 | a | L160526 |
| **pcf80** | old | 2.2632E+10 | 13235668964 | FT10b | F | T | 10 | FT10 | b | L160526 |
| **pcf76** | old | 2.2867E+10 | 13974587699 | FT8b | F | T | 8 | FT8 | b | L160526 |
| **pcf91** | old | 2.3215E+10 | 13798637662 | FT15c | F | T | 15 | FT15 | c | L160526 |
| **pcf21** | old | 2.3276E+10 | 14111350073 | FN8b | F | N | 8 | FN8 | b | L160526 |
| **pcf92** | old | 2.3327E+10 | 13858325588 | FT15d | F | T | 15 | FT15 | d | L160526 |
| **pcf50** | new | 2.3557E+10 | 12807009118 | FN36a | F | N | 36 | FN36 | a | L160530 |
| **pcf47** | new | 2.365E+10 | 11967196757 | FN34b | F | N | 34 | FN34 | b | L160530 |
| **pcf82** | old | 2.403E+10 | 13926924881 | FT11b | F | T | 11 | FT11 | b | L160526 |
| **pcf20** | old | 2.4037E+10 | 14520086962 | FN8a | F | N | 8 | FN8 | a | L160526 |
| **pcf100** | new | 2.4044E+10 | 13424517410 | FT33b | F | T | 33 | FT33 | b | L160529 |
| **pcf99** | new | 2.4128E+10 | 13514812329 | FT33a | F | T | 33 | FT33 | a | L160529 |
| **pcf81** | old | 2.4305E+10 | 14086354372 | FT11a | F | T | 11 | FT11 | a | L160526 |
| **pcf104** | new | 2.4402E+10 | 12831265087 | FT35b | F | T | 35 | FT35 | b | L160530 |
| **pcf103** | new | 2.4588E+10 | 13558991247 | FT35a | F | T | 35 | FT35 | a | L160530 |
| **pcf106** | new | 2.4613E+10 | 12228206196 | FT36b | F | T | 36 | FT36 | b | L160530 |
| **pcf52** | new | 2.4702E+10 | 12900362405 | FN37a | F | N | 37 | FN37 | a | L160530 |
| **pcf109** | new | 2.5214E+10 | 13770725148 | FT38a | F | T | 38 | FT38 | a | L160530 |
| **pcf157** | new | 2.5376E+10 | 15093982775 | PN34a | P | N | 34 | PN34 | a | L160530 |
| **pcf158** | new | 2.6005E+10 | 14676894860 | PN34b | P | N | 34 | PN34 | b | L160530 |
| **pcf48** | new | 2.6071E+10 | 13789627343 | FN35a | F | N | 35 | FN35 | a | L160530 |
| **pcf105** | new | 2.6243E+10 | 13742184910 | FT36a | F | T | 36 | FT36 | a | L160530 |
| **pcf110** | new | 2.6574E+10 | 13777156758 | FT38b | F | T | 38 | FT38 | b | L160530 |
| **pcf36** | old | 2.6673E+10 | 15522999281 | FN15c | F | N | 15 | FN15 | c | L160526 |
| **pcf37** | old | 2.6961E+10 | 15643813281 | FN15d | F | N | 15 | FN15 | d | L160526 |
| **pcf108** | new | 2.7714E+10 | 14059474140 | FT37b | F | T | 37 | FT37 | b | L160530 |
| **pcf45** | new | 2.7794E+10 | 14334047385 | FN33b | F | N | 33 | FN33 | b | L160529 |
| **pcf42** | new | 2.8089E+10 | 15579860949 | FN32a | F | N | 32 | FN32 | a | L160529 |
| **pcf44** | new | 2.8188E+10 | 14572088362 | FN33a | F | N | 33 | FN33 | a | L160529 |
| **pcf43** | new | 2.8269E+10 | 15763276246 | FN32b | F | N | 32 | FN32 | b | L160529 |
| **pcf219** | new | 2.8743E+10 | 16622795700 | PT36a | P | T | 36 | PT36 | a | L160530 |
| **pcf107** | new | 2.9031E+10 | 15313238354 | FT37a | F | T | 37 | FT37 | a | L160530 |
| **pcf55** | new | 2.9207E+10 | 14066768643 | FN38b | F | N | 38 | FN38 | b | L160530 |
| **pcf54** | new | 2.9382E+10 | 14847913252 | FN38a | F | N | 38 | FN38 | a | L160530 |
| **pcf97** | new | 3.3445E+10 | 18237981305 | FT32a | F | T | 32 | FT32 | a | L160529 |

**Supplementary Figure 7h. MS1 and MS2 TIC summary for the 224 SWATH files from the PCF data set sorted by ascending MS1 TIC sum value.** 80 samples with substantially low MS1 TIC values are shown in red text. 'pcf_id': the id of each swath file; 'sample_age': samples collected before 2010 are labeled "old", whereas samples collected after 2010 are labeled "new"; 'MS1_TIC': TIC sum value at MS1 level; 'MS2_TIC': TIC sum value at MS2 level; 'sample_name': name of each SWATH injection based on tissue type, storage method, patient and technical replicates; 'storage_type': P indicates FFPE while F means FF; 'patient_id': id of patient; 'sample_name2': sample name without distinguishing technical replicates; 'tech_rep': technical replicate id; 'MS_batch': SWATH acquisition batch.

**Supplementary Figure 8**

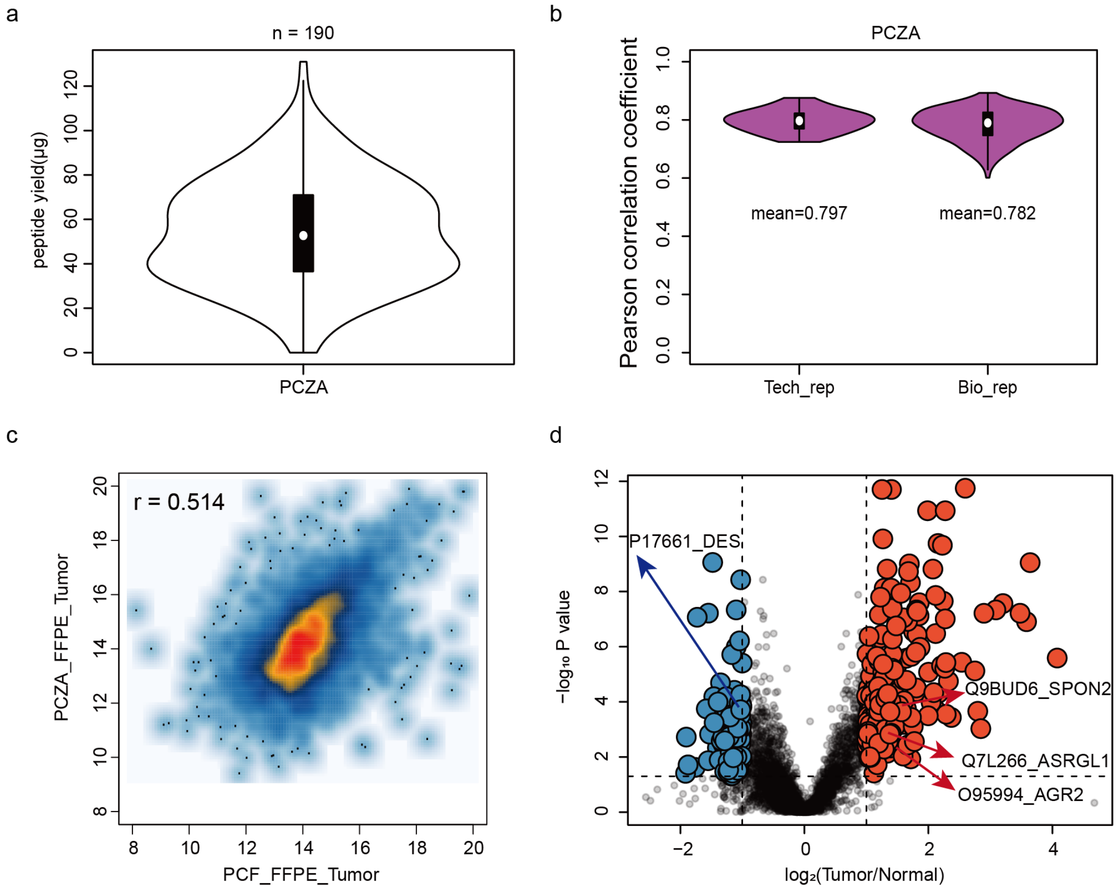


**Supplementary Figure 8. PCT-SWATH analysis of FFPE prostate cancer tissues from China (PCZA data set).** (a) Peptide yield per 1 mg FFPE tissue punch weight. (b) Technical and biological reproducibility of PCZA data set. Pearson correlation co-efficient based on protein expression was computed for technical and biological replicates respectively as measured by FFPE-PCT-SWATH. (c) Pearson correlation of PCF and PCZA FFPE tumor proteomes (r=0.514). (d) Volcano plot displaying the significantly regulated proteins between tumor and benign tissue samples. Red dots indicated upregulated proteins while blue ones indicated down regulated proteins.

### Supplementary Figure 9

Network 2 : PCZA_reg_prot_list_expression : PCZA_reg_prot_list : PCZA_reg_prot_list_expression

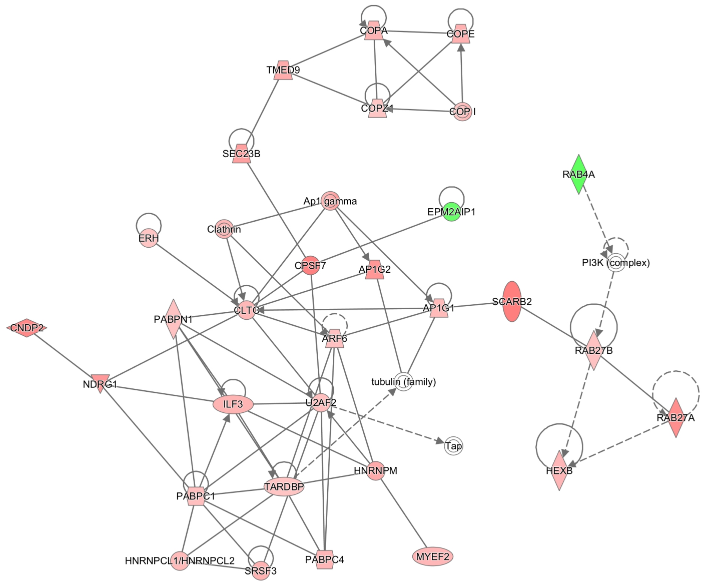


© 2000-2019 QIAGEN. All rights reserved.

Network 3 : PCZA_reg_prot_list_expression : PCZA_reg_prot_list : PCZA_reg_prot_list_expression

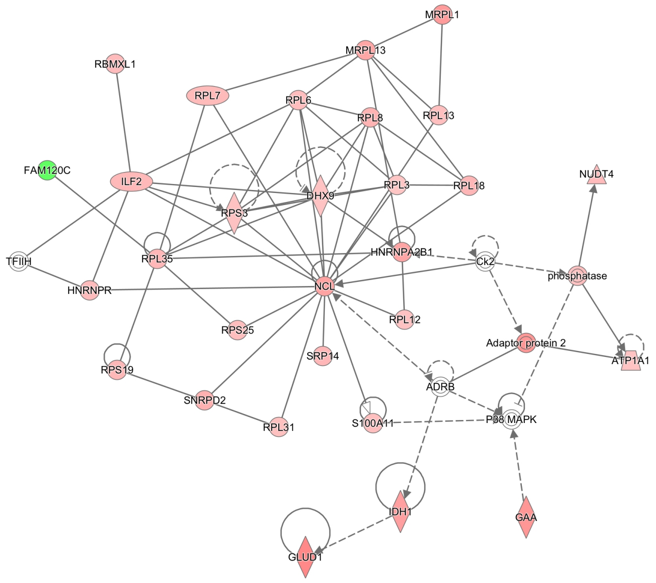


© 2000-2019 QIAGEN. All rights reserved.

Network 5 : PCZA_reg_prot_list_expression : PCZA_reg_prot_list : PCZA_reg_prot_list_expression

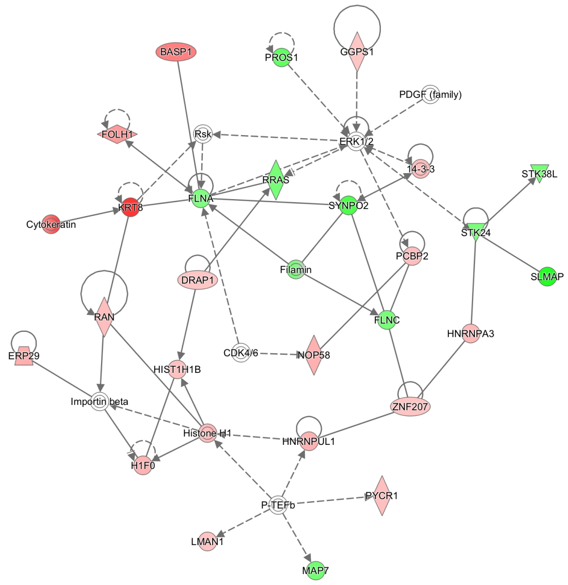


© 2000-2019 QIAGEN. All rights reserved.

Network 6 : PCZA_reg_prot_list_expression : PCZA_reg_prot_list : PCZA_reg_prot_list_expression

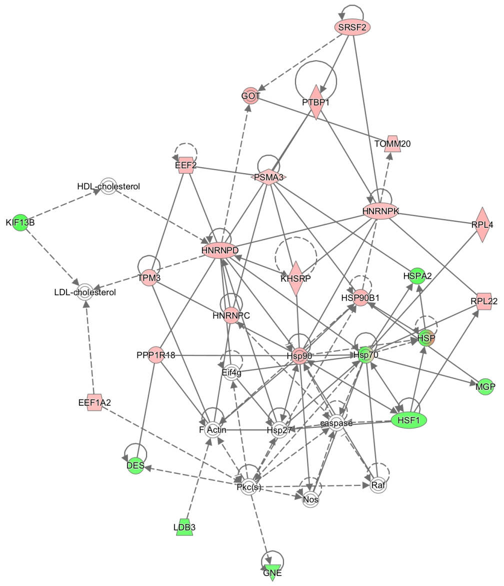


© 2000-2019 QIAGEN. All rights reserved.

Network 7 : PCZA_reg_prot_list_expression : PCZA_reg_prot_list : PCZA_reg_prot_list_expression

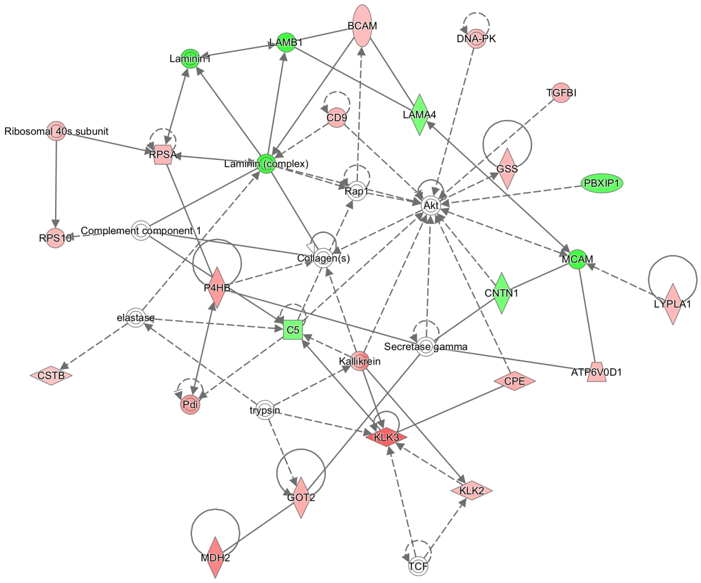


© 2000-2019 QIAGEN. All rights reserved.

Network 9 : PCZA_reg_prot_list_expression : PCZA_reg_prot_list : PCZA_reg_prot_list_expression

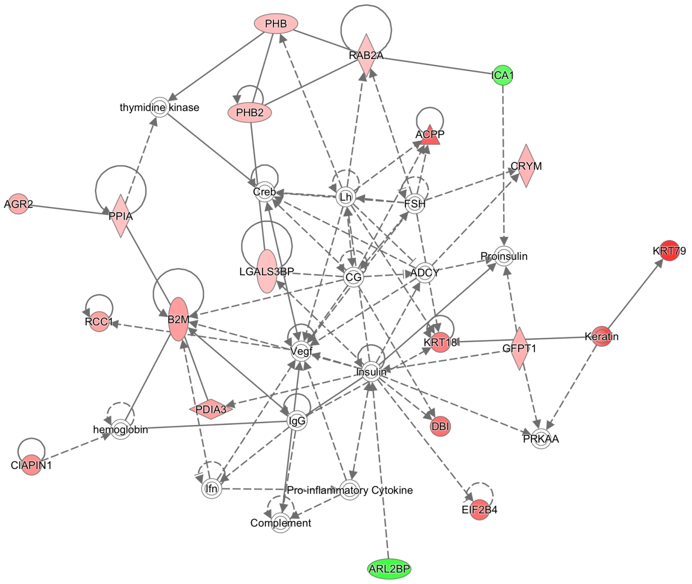


© 2000-2019 QIAGEN. All rights reserved.

Network 11 : PCZA_reg_prot_list_expression : PCZA_reg_prot_list : PCZA_reg_prot_list_expression

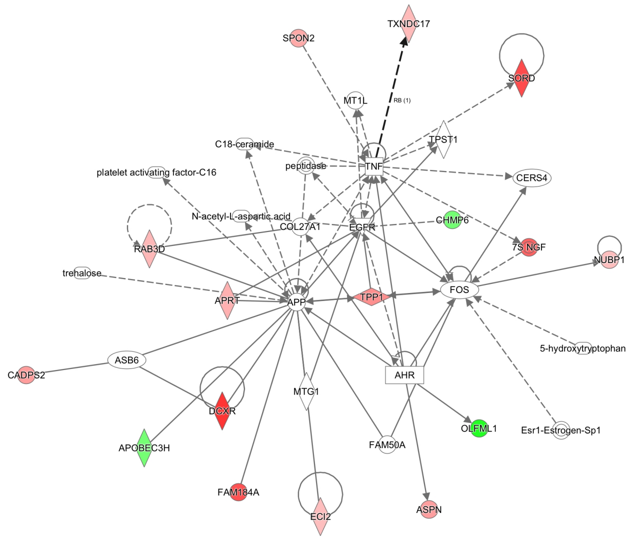


© 2000-2019 QIAGEN. All rights reserved.

Network 13 : PCZA_reg_prot_list_expression : PCZA_reg_prot_list : PCZA_reg_prot_list_expression

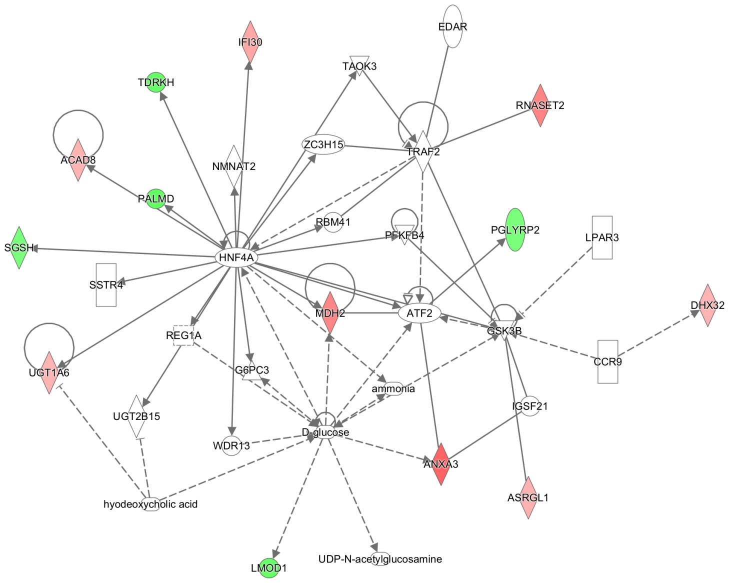


© 2000-2019 QIAGEN. All rights reserved.

Network 15 : PCZA_reg_prot_list_expression : PCZA_reg_prot_list : PCZA_reg_prot_list_expression

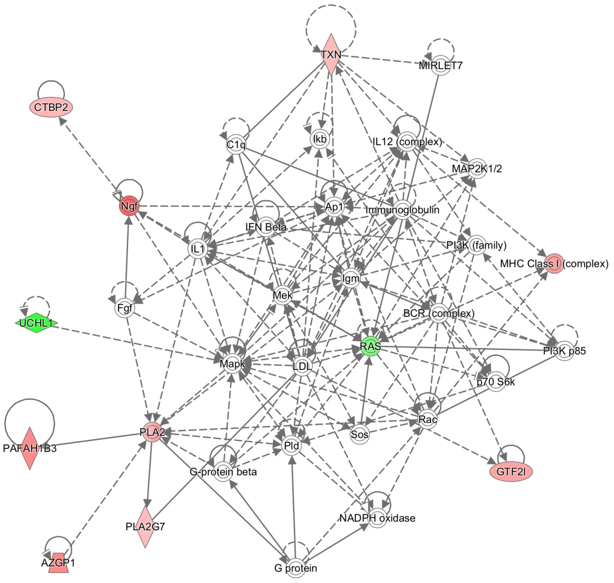


© 2000-2019 QIAGEN. All rights reserved.

**Supplementary Figure 9. Selected nine networks enriched by IPA analysis.** The eleven proteins KLK3, SORD, UCHL1, FLNA, TPP1, HNRNPA2B1, MDH2, AGR2, SPON2, ASRGL1, and DES were involved in nine networks which are numbered with 2, 3, 5, 6, 7, 9, 11, 13, 15, respectively. Pink dots represent proteins which were upregulated in tumor tissues while green ones were downregulated.

#### Supplementary Figure 10

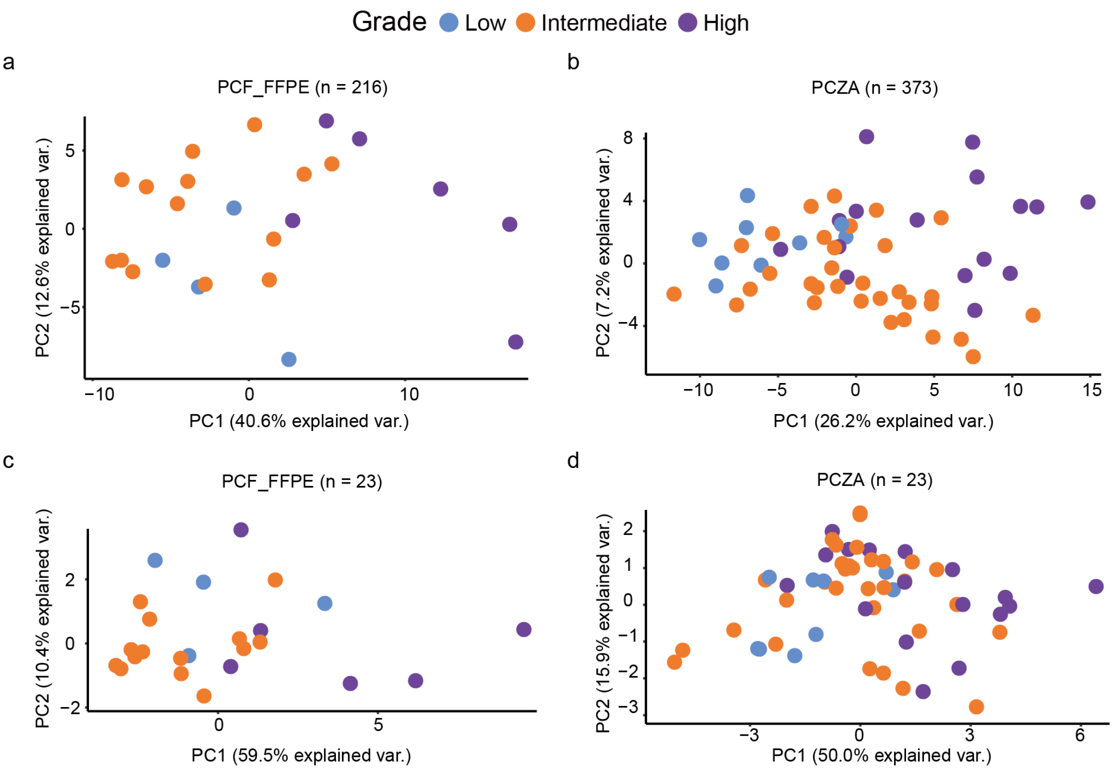


**Supplementary Figure 10. Using a panel of featured proteins generated by ANOVA analysis in pairwise among three disease stages, low grade, intermediate grade, and high grade, to distinguish different stages. (a)** 216 selected proteins were employed to distinguish low and high grades in PCF cohort. **(b)** 373 proteins were employed to distinguish low and high grades in PCZA cohort. **(c)** 23 proteins in common were employed to distinguish low and high grades in PCF cohort. **(d)** 23 proteins in common were employed to distinguish low and high grades in PCZA cohort.

#### Supplementary Figure 11

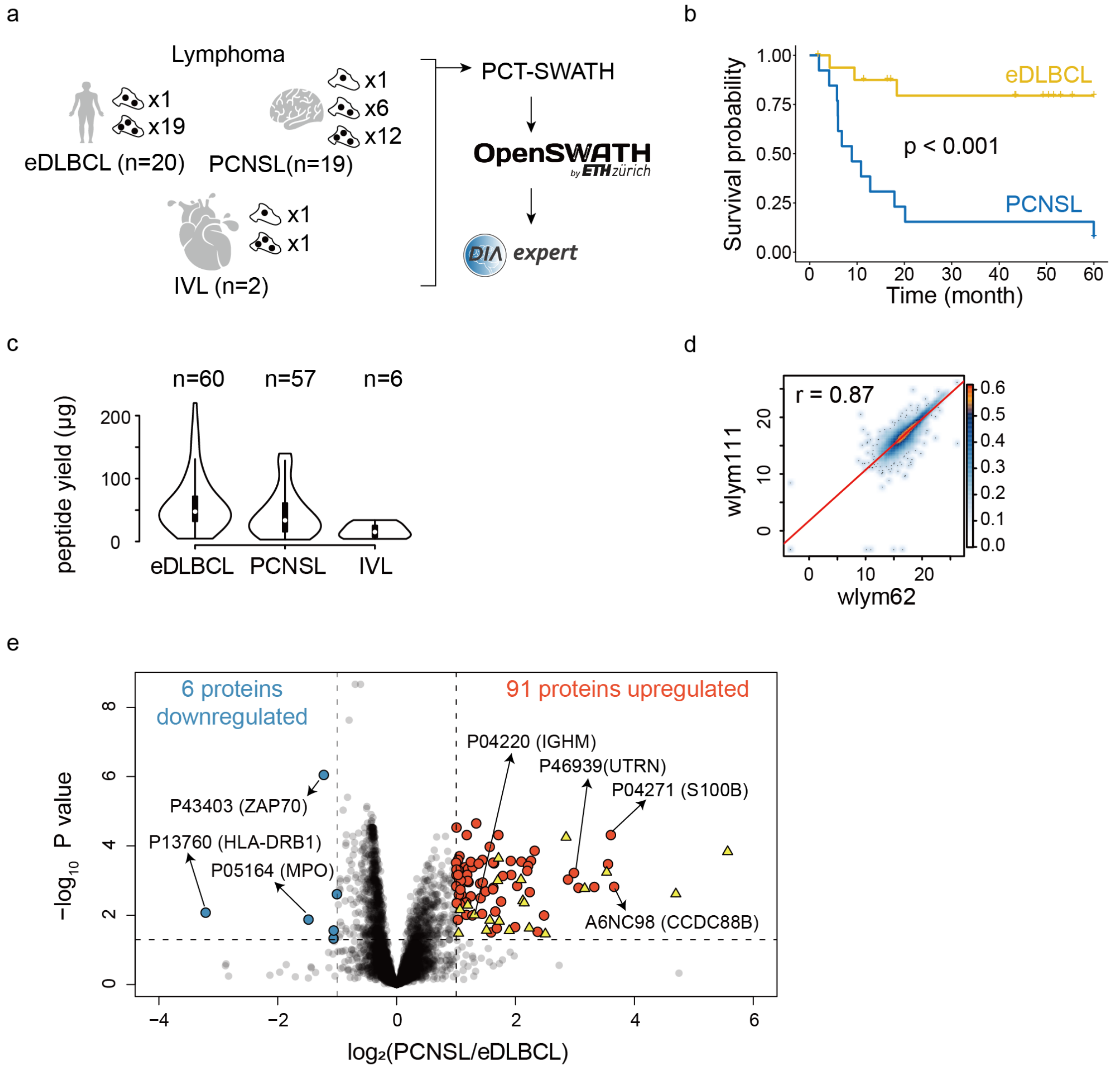


**Supplementary Figure 11. PCT-SWATH analysis of FFPE diffuse large B-cell lymphoma tissues.** (**a**) Study design. 20 eDLBCL, 19 PCNSL and 2 IVL patients were recruited. From resected tissue of each patient two or three FFPE punches were sampled and analyzed by PCT-SWATH. (**b**) Survival analysis of eDLBCL and PCNSL patients up to 60 months. (**c**) Peptide yield per 1 mg FFPE tissue punch weight. 'n' means the number of tissue punches. (**d**) Technical reproducibility from a randomly selected eDLBCL sample. (**e**) Volcano plot displaying the significantly regulated proteins between PCNSL and eDLBCL samples. Yellow triangles indicated brain proteins that are presumably contaminants in the lymphoma sample. IVL samples were not analyzed in this regard due to small sample size.

#### Supplementary Figure 12

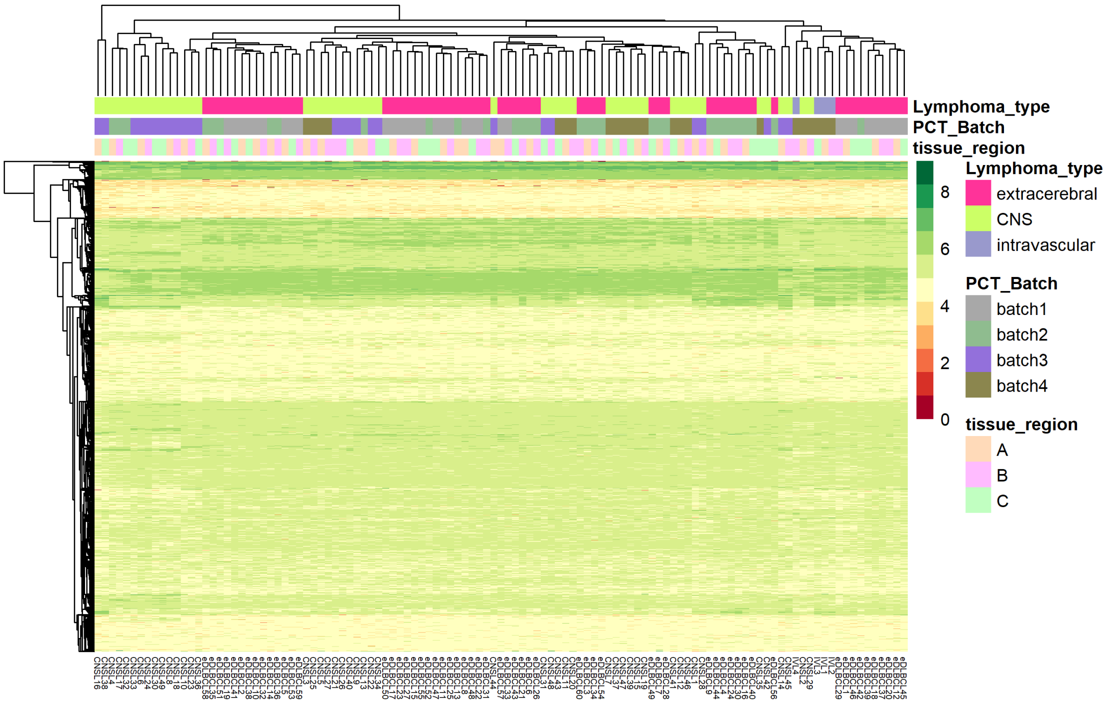


**Supplementary Figure 12. Unsupervised clustering of 113 lymphoma samples.** Unsupervised clustering of 113 Diffuse Large B cell Lymphoma samples based on 5769 SwissProt proteins.

#### Supplementary Fig. 13

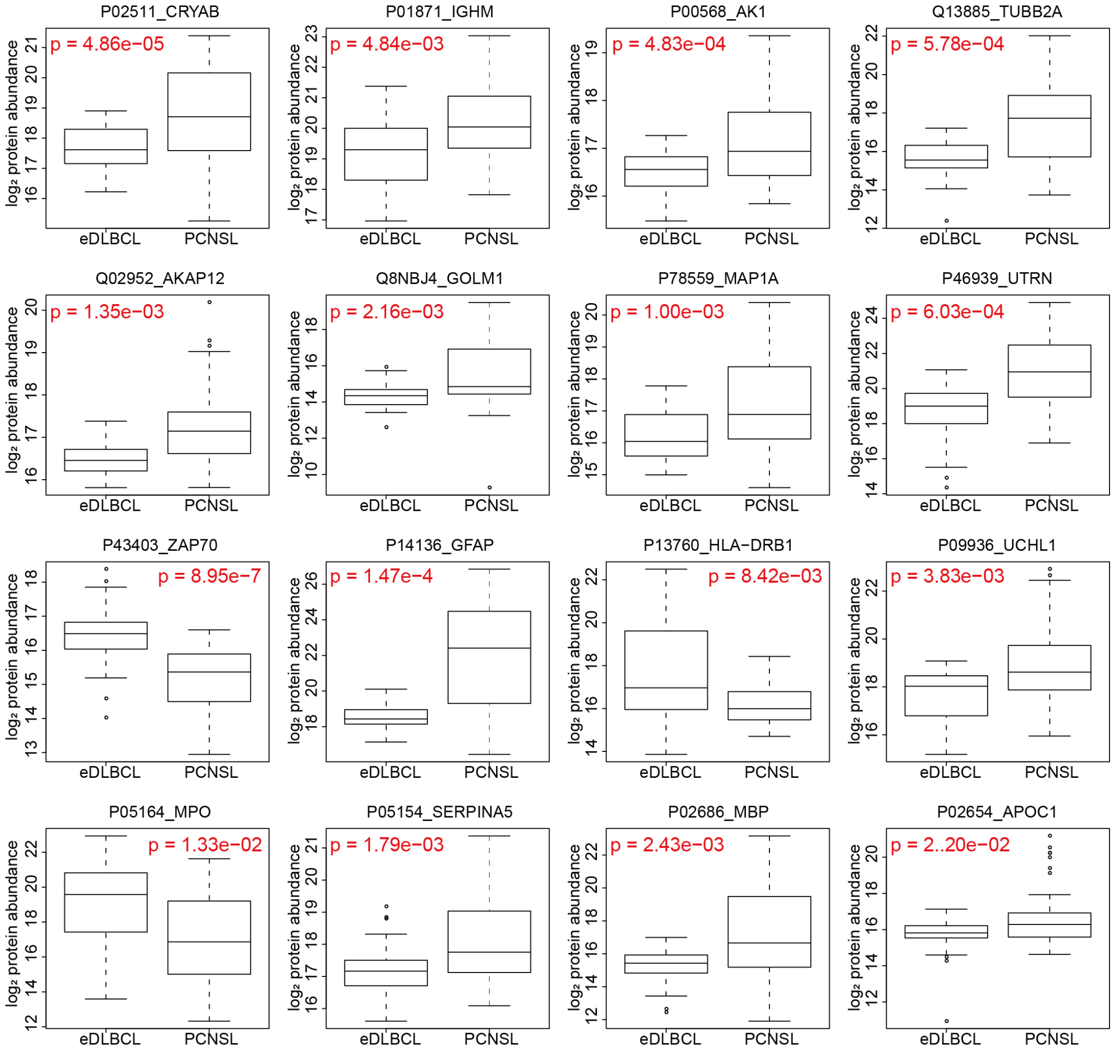


**Supplementary Figure 13. Abundance of the sixteen proteins in eDLBCL/PCNSL samples in WLYM cohort.** They were also detected in ZLYM eDLBCL cohort.

**Supplementary Fig. 14**

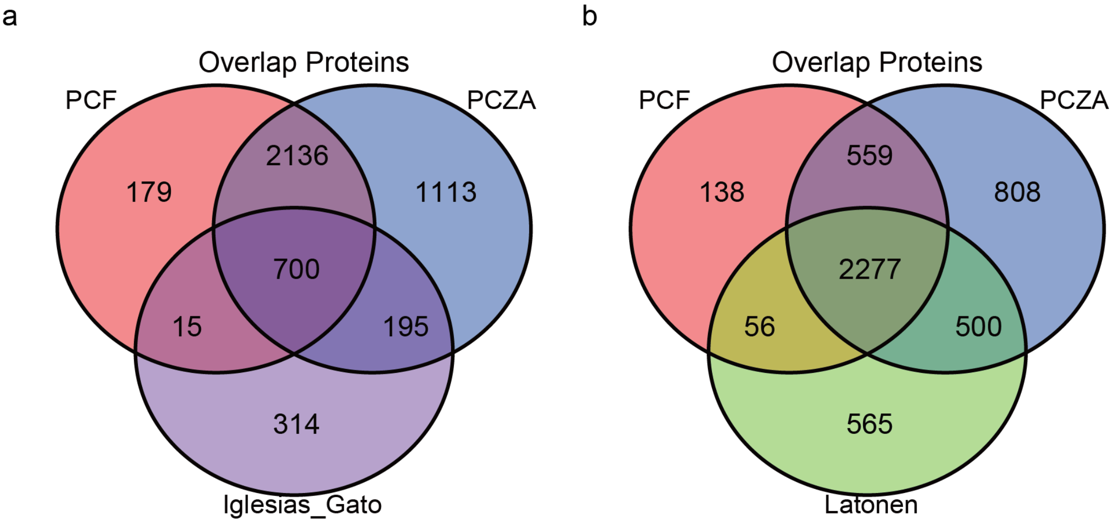


**Supplementary Figure 14**. Venn diagram of overlapped proteins from PCF, PCZA, Iglesias-Gato (78) and Latonen (61) proteomes. (**a**) 700 common proteins in PCF, PCZA and Iglesias-Gato proteomes. (**b**) 2277 common proteins in PCF, PCZA and Latonen proteomes.
